# Decoding Substrate Specificity in a Promiscuous Biocatalyst by Enzyme Proximity Sequencing

**DOI:** 10.1101/2025.07.10.664162

**Authors:** Rosario Vanella, Sean Boult, Christoph Küng, Michael A. Nash

## Abstract

Substrate specificity is a defining feature of enzyme function, but its molecular underpinnings remain difficult to decode and engineer. Here, we leveraged enzyme proximity sequencing (EP-Seq) to systematically map how single-point and combinatorial mutations reshape the substrate preferences of D-amino acid oxidase (DAOx) from *Rhodotorula gracilis*, a model promiscuous enzyme. We generated ∼40,000 sequence–phenotype pairs, enabling us to profile the activities of ∼6,500 unique DAOx variants against five D-amino acid substrates with distinct physicochemical properties. Our analysis revealed that substrate-specific mutations are distributed throughout the enzyme structure. Mutations near the active site drive strong specificity shifts but also incur catalytic penalties, while distal mutations subtly rewire intramolecular contacts in order to modulate specificity with minimal loss of activity. We identified and validated positional hotspots that act allosterically to influence specificity, and characterized key variants that acquired exclusive substrate specificity or exhibited up to 230-fold changes in substrate preference. Combining mutations with complementary effects further sharpened substrate discrimination, enabling rational design of highly selective biocatalysts. This work provides a powerful framework for decoding enzyme specificity and provides unique foundational datasets to advance AI-guided enzyme engineering.

## Introduction

Tailoring substrate specificity in enzymes is crucial for biocatalyst engineering, therapeutic enzyme design, and synthetic biology. Despite major progress in structural and computational enzymology, our understanding of how enzymes discriminate between structurally similar substrates remains incomplete, particularly in enzymes that exhibit broad promiscuity. Substrate specificity was for many years considered a static property^1^, but is now appreciated as an emergent feature shaped by an interplay of active-site geometry, conformational dynamics, and long-range intramolecular interaction networks, all under complex evolutionary constraints^2–4^.

Substrate specificity manifests as a relative property, enabling enzymes to preferentially convert their primary substrate relative to structurally similar alternatives, while maintaining sufficient reaction flux in the crowded cellular environment^3,5,6^. However, the molecular principles governing substrate discrimination and its co-evolution with other enzyme properties (e.g., overall catalytic activity and folding stability) remain poorly understood. This knowledge gap limits our ability to predict how mutations will affect enzyme performance and impedes the rational engineering of enzymes for diverse applications.

Deep mutational scanning (DMS) has revolutionized the study of protein function by enabling high-throughput mapping of sequence-fitness relationships^7–9^. If successfully applied to the study of enzyme specificity, DMS could have significant potential to illuminate how sequence encodes discrimination between structurally similar substrates. However, applying DMS to enzymes presents unique challenges. DMS workflows applied to enzymes must directly link catalytic activity to a measurable cellular phenotype. This requirement has largely limited prior work to enzymes whose activity can be directly linked with cell growth or survival^10–12^. Prior DMS work on substrate specificity has therefore focused on antibiotic resistant β-lactamase^13–15^ and on an amide hydrolase^16^ whose function generates a nitrogen source required for cell proliferation.

These prior studies demonstrate that each substrate shapes the enzyme’s fitness landscape in a distinct way, and offer preliminary insights into molecular determinants of specificity. However, this line of research has been limited by reliance on intracellular reactions and growth-based selections, where reaction conditions may be poorly controlled, substrate concentrations cannot be precisely set, and the observed phenotypic fitness may conflate the influence of mutations on catalytic activity with their impact on enzyme abundance (i.e., expression level) or folding stability.

To overcome these challenges, we recently developed enzyme proximity sequencing (EP-Seq), a DMS workflow that leverages a high-throughput single-cell enzyme activity assay based on radical-mediated fluorescent proximity labeling^17–21^. EP-Seq enables quantification of enzyme activity for cells displaying an enzyme variant on their surface, where catalytic turnover can be linked, directly or indirectly via a cascade reaction, to the production of hydrogen peroxide (H_2_O_2_). We previously validated EP-Seq by mapping the mutational landscape of an oxidoreductase^17^, showing that the method can disentangle the effects of mutations on expression level, folding stability, and catalytic activity in a massively parallel and quantitative manner.

In the present work, we extend the use of EP-Seq to examine how mutations affect substrate selectivity in a D-amino acid oxidase from *Rhodotorula gracilis*, a naturally promiscuous enzyme with diverse industrial and biomedical applications^22–25^. By displaying a comprehensive library of single-substitution enzyme variants on the surface of yeast cells, we measured the activity of thousands of variants against five substrates with distinct biophysical properties. This setup allowed precise control over reaction conditions and facilitated direct comparison of fitness scores across datasets.

By analyzing the fitness landscape of DAOx on five substrates, we identified hundreds of single amino acid substitutions that differentially modulate enzyme performance, revealing how mutations that control substrate preference are distributed across the entire protein structure. Mutations near the active site produced large specificity shifts but often incurred catalytic penalties, while distal substitutions more subtly fine-tuned substrate preferences with minimal loss of activity. This highlights distinct mechanisms for achieving specificity depending on the extent of functional cost that can be tolerated during evolution. We further developed a substrate specificity score, and validated this statistical metric using 140 single-clone measurements of enzymatic discrimination factors. This high-throughput DMS-based phenotypic metric enables rapid and quantitative assessment of enzyme selectivity between substrate pairs. Finally, we showed that combining individually beneficial mutations can synergistically sharpen substrate specificity, offering a scalable route to engineering highly selective biocatalysts. Together, these findings establish EP-Seq as a robust platform for decoding and engineering enzyme specificity at scale.

## Results

### EP-Seq workflow applied to study substrate specific effects of mutations

To systematically map how mutations affect substrate specificity in *Rhodotorula gracilis* D-amino acid oxidase (DAOx) we applied EP-Seq^17^, a high-throughput platform that links enzyme activity to a fluorescence readout at the single-cell level. In this workflow, a library of enzyme variants fused to the Aga2 anchor protein is displayed on the surface of *Saccharomyces cerevisiae*. Enzymatic turnover is coupled to production of H_2_O_2_, which in the presence of horseradish peroxidase (HRP) initiates a tyramide-based radical labeling reaction that links fluorophores to the cell surface. The resulting cellular fluorescence intensity correlates with the catalytic turnover of the variant borne by that particular cell. Following labeling, cells are sorted by fluorescence activated cell sorting (FACS) into bins, and next-generation sequencing (NGS) analysis of the DNA barcodes reveals the identities of the enzyme variants in each bin. Fitness scores for each variant are then computed by comparing their bin distributions to that of the wild-type enzyme, allowing quantitative measurements of activity. In a parallel experiment, the expression levels of all variants in the library are monitored by fluorescently labelling a C-terminal epitope tag (e.g., 6×His tag) on the displayed enzyme construct, and again quantifying the expression fitness by FACS and NGS. By decoupling activity and expression measurements, EP-Seq enables the construction of orthogonal fitness landscapes for catalytic activity and display levels.

We applied EP-Seq to evaluate the effects of single-point mutations on the catalytic activity of DAOx across five distinct D-amino acid substrates (**Fig. 1A**). The substrate panel included three hydrophobic amino acids (D-alanine (D-Ala), D-methionine (D-Met), and D-phenylalanine (D-Phe)) with progressively larger side chains (methyl, S-methyl thioether, and phenyl, respectively). Additionally, we included two hydrophilic substrates (D-asparagine (D-Asn), and D-glutamine (D-Gln)) bearing polar amide groups on two– and three-carbon side chains, respectively.

**Figure 1.**
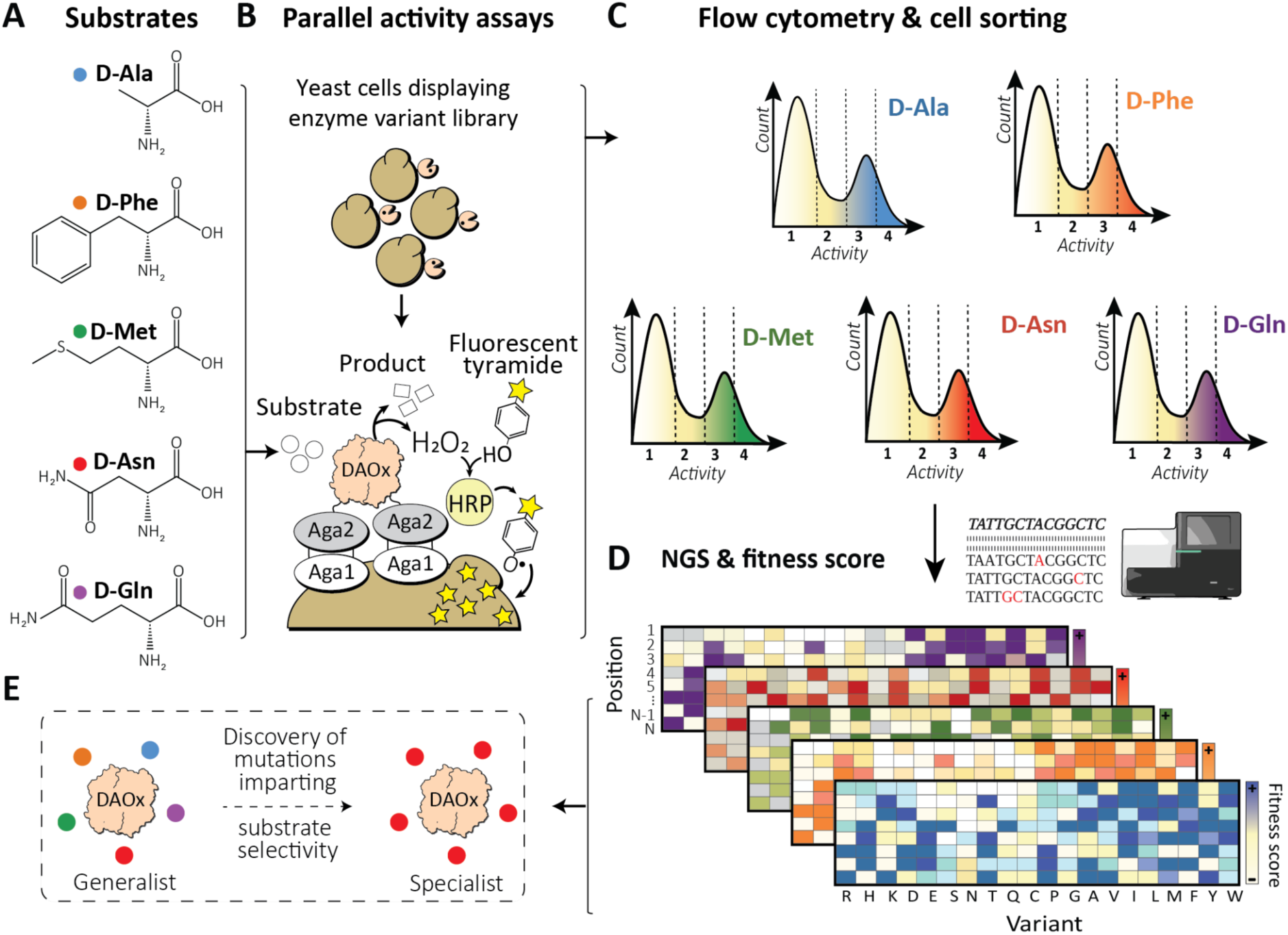
EP-Seq workflow for mapping substrate specificity in the promiscuous enzyme D-amino acid oxidase (DAOx). (**A**) Five D-amino acid substrates with distinct physicochemical properties were selected to profile the activity of a comprehensive DAOx variant library. (**B**) Yeast cells display DAOx variants on their surface. Enzyme activity is coupled to H_2_O_2_ production, which initiates a tyramide-based labeling reaction catalyzed by horseradish peroxidase (HRP). This covalently attaches fluorescent radicals onto cells displaying active variants, linking catalytic activity to fluorescence intensity. (**C**) Each substrate assay is analyzed by flow cytometry and fluorescence-activated cell sorting (FACS), partitioning cells into four activity-based bins. (**D**) Barcode sequencing of the sorted populations reveals the identity and abundance of variants in each bin, allowing computation of a fitness score that quantifies activity on each substrate relative to WT. (**E**) Comparative analysis across substrates identifies specificity-determining mutations, reveals structural hotspots for engineering selective variants, and provides insights into the biophysical trade-offs between activity, specificity and stability.

We first cloned the wild type DAOx (WT-DAOx) gene in-frame with Aga2, and confirmed that the surface-displayed fusion protein on *S. cerevisiae* strain EBY100 retained activity and substrate preferences comparable to the soluble enzyme. We found that the hydrophobic substrates were processed with higher affinity than the polar ones (**Note S1, Fig. S1, Table S1**). For the EP-Seq assays, each substrate was assayed at its experimentally determined *K*_M_ concentration. Time-course optimization of the tyramide reaction showed that a 60-minute reaction time yielded optimal labeling for the hydrophobic substrates, while 120 minutes were required for the less reactive polar substrates. These conditions ensured comparable signal intensities and dynamic ranges across all five substrates (**Note S2, Fig. S2**).

We next transformed EBY100 yeast cells with a site-saturation mutagenesis library of DAOx variants, covering 94.4% of all possible single amino acid substitutions (6,530 of 6,916 possible missense variants; **Fig. 1B**). After the tyramide labeling assay, cells were sorted by fluorescence intensity. A low fluorescence gate captured cells either not displaying the enzyme or bearing inactive variants, while the remaining cells were equally divided into three bins of increasing signal (**Fig. 1C, Fig. S3**). Barcode sequencing of each bin was then used to reveal the identity and frequency of the variants, from which we computed an activity fitness score for each variant relative to WT (**Fig. 1D-E, Table S2**, **Eqs. 1-3**).

### Validation of the EP-Seq workflow and analysis of fitness scores

To assess the reproducibility of the workflow, we calculated Pearson correlation coefficients (*r* values) between fitness scores obtained from two biological replicates. This analysis included single-mutant variants represented by at least 10 cells per replicate at the screening stage. Across all five substrates, Pearson’s *r* values ranged from 0.94 to 0.97, indicating high reproducibility of the assay (**Fig. 2A**).

**Figure 2.**
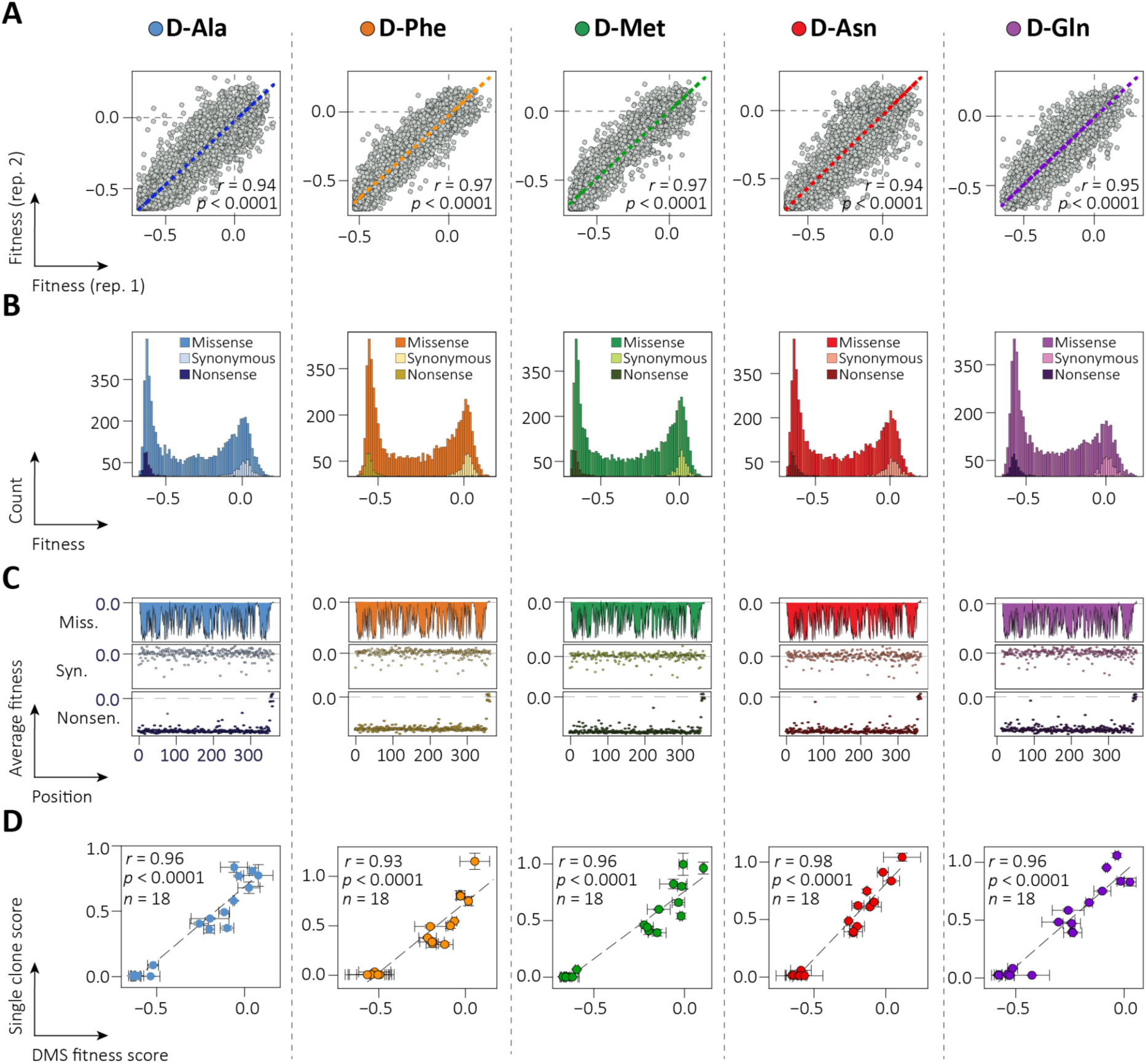
Validation of EP-Seq fitness scores quantifying DAOx activity across five substrates. (**A**) Scatter plots and linear regression of activity scores calculated from two biological replicates for each of the five substrates. (**B**) Distribution of activity fitness scores for missense, synonymous, and nonsense mutations across all variants and substrates. (**C**) Average fitness effect per sequence position for missense (top), synonymous (middle) and nonsense (bottom) mutations. (**D**) Correlation between DMS-derived fitness scores and catalytic activity measured independently in monogenic yeast culture for 18 single DAOx variants. Dashed line: linear regression; *r*: Pearson correlation coefficients, n: number of datapoints, *p*: p-values two-tailed.

Within each screen, we computed a consensus score for each DAOx variant (**Methods, Eq. 4**), and examined the distributions of scores for missense, nonsense and synonymous mutations (**Fig. 2B-C**). Missense mutations were generally associated with reduced catalytic activity (average score = –0.31 ± 0.05; n = 5,821 variants in all five datasets) (**Fig. 2B-C, Table S3**). Nonsense mutations had strongly negative effect on activity (average score –0.58 ± 0.05; n = 316 in all five datasets), indicating that loss of the C-terminal region beginning anywhere from position 1 to position 354 had a deleterious effect on the catalytic activity of the enzyme, regardless of the substrate used (**Table S3**). In contrast, variants carrying premature stop codons beyond position 356, which resulted in loss of only the last ∼10 amino acids, exhibited fitness scores comparable to WT (average score –0.01 ± 0.03; n = 9 variants in all five datasets) (**Fig. 2C**), suggesting that this C-terminal region is largely unnecessary for catalytic function, regardless of the substrate. Synonymous mutations did not substantially alter fitness, with scores similar to WT (avg score = 0.00 ± 0.0035; n = 281 in all five datasets) (**Fig. 2B-C, Table S3**).

To validate the fitness scores obtained from EP-Seq, we used an Amplex Red assay to measure the catalytic activity of 18 yeast-display single-mutant DAOx variants on each of the five substrates. Initial reaction rates were normalized to that of the WT enzyme assayed in parallel and compared to the corresponding DMS-derived fitness scores. The two measurements were highly consistent, with Pearson’s *r* values ranging from 0.93 to 0.98 across all tested conditions (**Fig. 2D**).

### Comparative analysis of EP-Seq scores across five DAOx substrates

We then combined the five high-quality EP-Seq datasets. The resulting data comprised 6,418 unique variants, including 5,821 missense, 316 nonsense, and 281 synonymous mutations. To account for differences in enzyme display (i.e. expression) levels, activity scores were normalized to expression scores obtained from our previously published dataset^17^. We further only considered variants with expression levels higher than the mean expression of nonsense variants plus one standard deviation. Synonymous variants were used to define a reference range for WT-like behavior (average score across all datasets: 1.00 ± 0.037). Nonsense and synonymous mutations were subsequently excluded, leaving 5,800 missense variants (83.9% of the total) shared across all five substrate screens that were used for comparative downstream analysis.

To qualitatively compare the effects of mutations, we visualized the fitness scores using heatmaps, which revealed broadly similar activity landscapes across all five screens (**Fig. S4**). Quantitative analysis further confirmed strong concordance in these datasets, with Pearson’s *r* values ranging from 0.93 to 0.96 in pairwise comparisons between different substrates (**Fig. S5**). The weakest correlation (*r* = 0.93) was observed between D-Ala and D-Gln (i.e. the smallest hydrophobic and the largest polar substrates, respectively). In contrast, the strongest correlations (*r* = 0.96) were found between D-Met and each of D-Phe, D-Ala, D-Asn, highlighting the intermediate size and chemical features of D-Met in comparison to the other substrates tested.

We attribute the high level of agreement to the inherently promiscuous and generalist nature of *R. gracilis* DAOx, a trait that is conserved in more evolutionarily recent homologs such as human DAOx. Additionally, our use of standardized assay conditions (i.e. *K*_M_-matched substrate concentrations and optimized timepoints) facilitated a consistent dynamic range across all substrate screens, despite significant substrate-dependent differences in catalytic efficiency. This minimized potential detection biases and enhanced scoring accuracy, particularly for low-activity variants where substrate-specific effects are most likely to occur due to functional trade-offs in enzyme evolution ^3,13,17^.

### Identification of specificity-determining positions in DAOx

To identify mutations that alter substrate preference, we computed z-scores for each missense variant across all pairwise combinations of the five substrate datasets. These z-scores quantify how much a variant’s fitness difference between two substrates deviates from the average difference observed across all mutations between two substrates. A positive z-score indicates enhanced activity on one substrate relative to its paired counterpart, while a negative z-score indicates the opposite (**Fig. S6**). We refer to this metric as the substrate specificity score (Sp-score), which captures both the magnitude and direction of a mutation’s impact on substrate specificity (**Methods, Eq. 5**).

To define a significance threshold, we used synonymous mutations as an internal control population to estimate background noise. Mutations with |Sp-score| ≥ 3, corresponding to three standard deviations from the mean, were classified as substrate-specific mutations. This threshold excluded >99% of synonymous variants and retained 601 missense mutations (332 unique variants) that were then classified as substrate-specific (**Fig. 3A-B, Fig. S6**). These mutations were broadly distributed throughout the DAOx sequence, affecting 166 of 364 positions (∼45%) (**Fig. 3A**).

**Figure 3.**
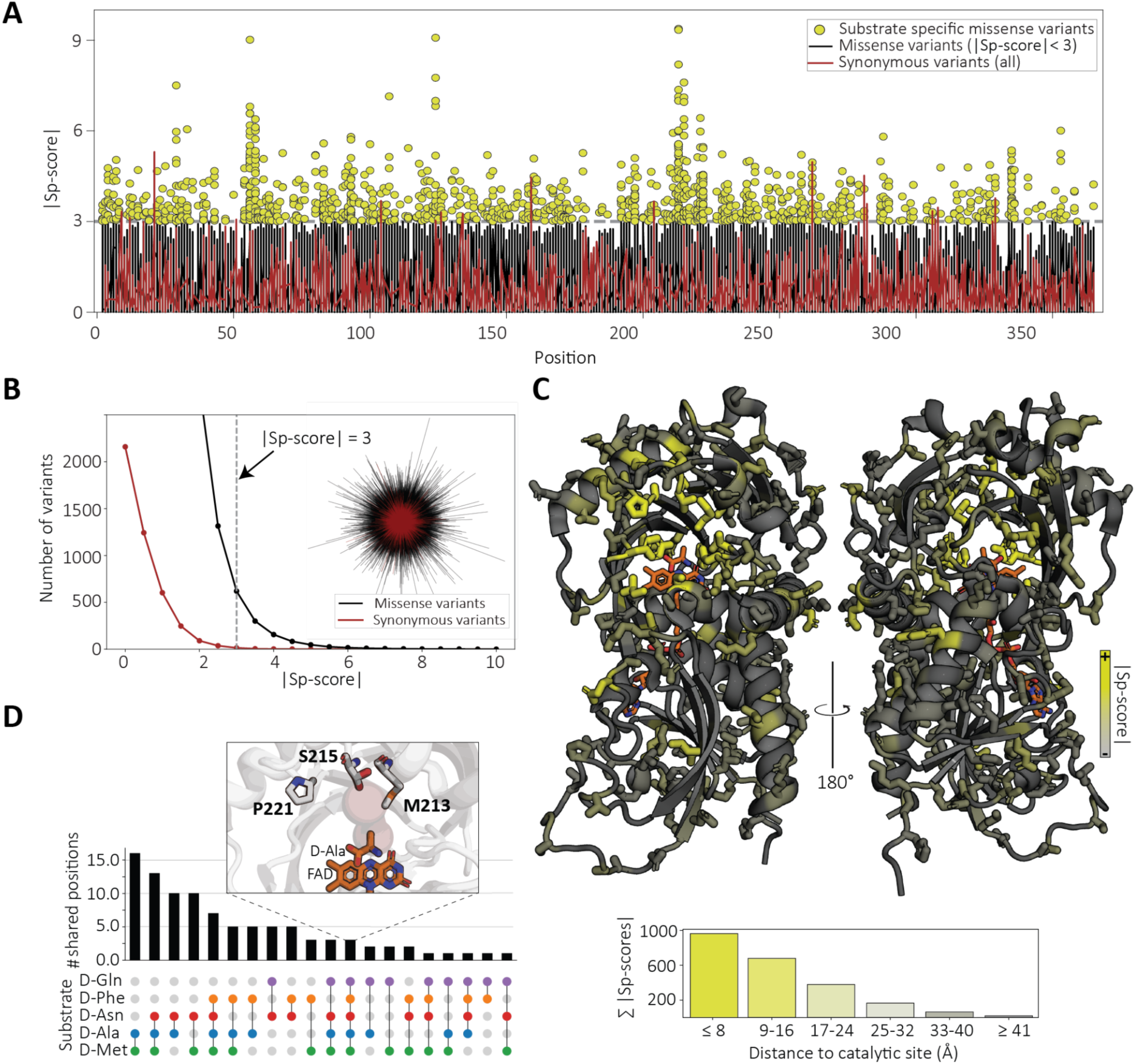
Identification and structural mapping of DAOx mutations conferring substrate specificity. (**A**) Scatter plot of absolute specificity-scores for missense and synonymous variants across DAOx positions. Missense variants with |Sp-score| ≥ 3 are shown as yellow dots. (**B**) Relationship between the number of variants and their corresponding absolute z-scores. A threshold of |Sp-score| ≥ 3 was selected to include <1% of synonymous mutations. Inset: starburst plot comparing z-score distributions of missense and synonymous variants, illustrating the clustering of synonymous mutations near zero Sp-score. Sp-score increases radially outward from the center. (**C**) DAOx crystal structure (PDB ID: 1C0P) with substrate-specific positions highlighted as sticks. Color intensity (grey to yellow) represents the cumulative absolute Sp-score per position, indicating how frequently each site contributes to substrate specificity. Bottom: bar plot showing the relationship between the distance of substrate-specific mutations from the catalytic site and their cumulative absolute specificity scores. (**D**) UpSet plot showing unique and shared DAOx positions with substrate-specific effects across the five tested substrates. Inset: zoom-in on positions 213, 215, and 221, which alter substrate specificity across all five substrates depending on the substituting residue (**see Fig. S7**).

When mapped onto the 3D structure, substrate-specific mutations appeared throughout the enzyme, with an average distance of 18–24 Å from the catalytic center depending on the substrate. However, mutations ≤ 8 Å from the active site tended to show the strongest effects (i.e., highest |Sp-score|), highlighting the important contribution of active-site and first shell residues to specificity (**Fig. 3C**).

While most specificity-determining positions were unique to one or two substrates, a small set showed broader effects. We found three positions (residues 213, 215, and 221) that exhibited differential substrate-specific effects across all five tested substrates, depending on the identity of the introduced amino (**Fig. 3D, Fig. S7**). The greatest overlap in substrate-specific positions was observed between D-Ala and D-Met (n = 15), suggesting a convergent mutational pattern shaped by their similar physicochemical profiles. In contrast, D-Phe and D-Gln shared only one specificity-associated site despite being similarly bulky. This limited overlap highlights the influence of chemical composition over steric similarity in shaping the specificity landscape (**Fig. 3D**).

### Validation and global analysis of substrate specificity scores

To validate the EP-Seq-derived substrate specificity scores (Sp-scores), we performed kinetic measurements on 17 single-mutant DAOx variants exhibiting high Sp-scores towards at least one of the five substrates. Each variant was displayed on yeast, and Michaelis-Menten kinetics were measured on all five substrates. For each variant and substrate pair, we then calculated discrimination factors (D-factors). The D-factor, defined as (*k*_cat_/*K*_M_)_sub1_ / (*k*_cat_/*K*_M_)_sub2_ (**Methods, Eq.8**) provides a standardized metric of specificity, representing catalytic efficiency on a cognate substrate divided by that on a non-cognate substrate^3,6^. We then compared Sp-scores with D-factors across 140 variant-substrate combinations. The two measures were strongly correlated (Pearson’s *r* = 0.76, *p* < 0.0001, *n* = 140, **Fig. 4A**), confirming that the EP-Seq-derived SP-scores accurately reflect substrate-selective catalytic behavior.

**Figure 4.**
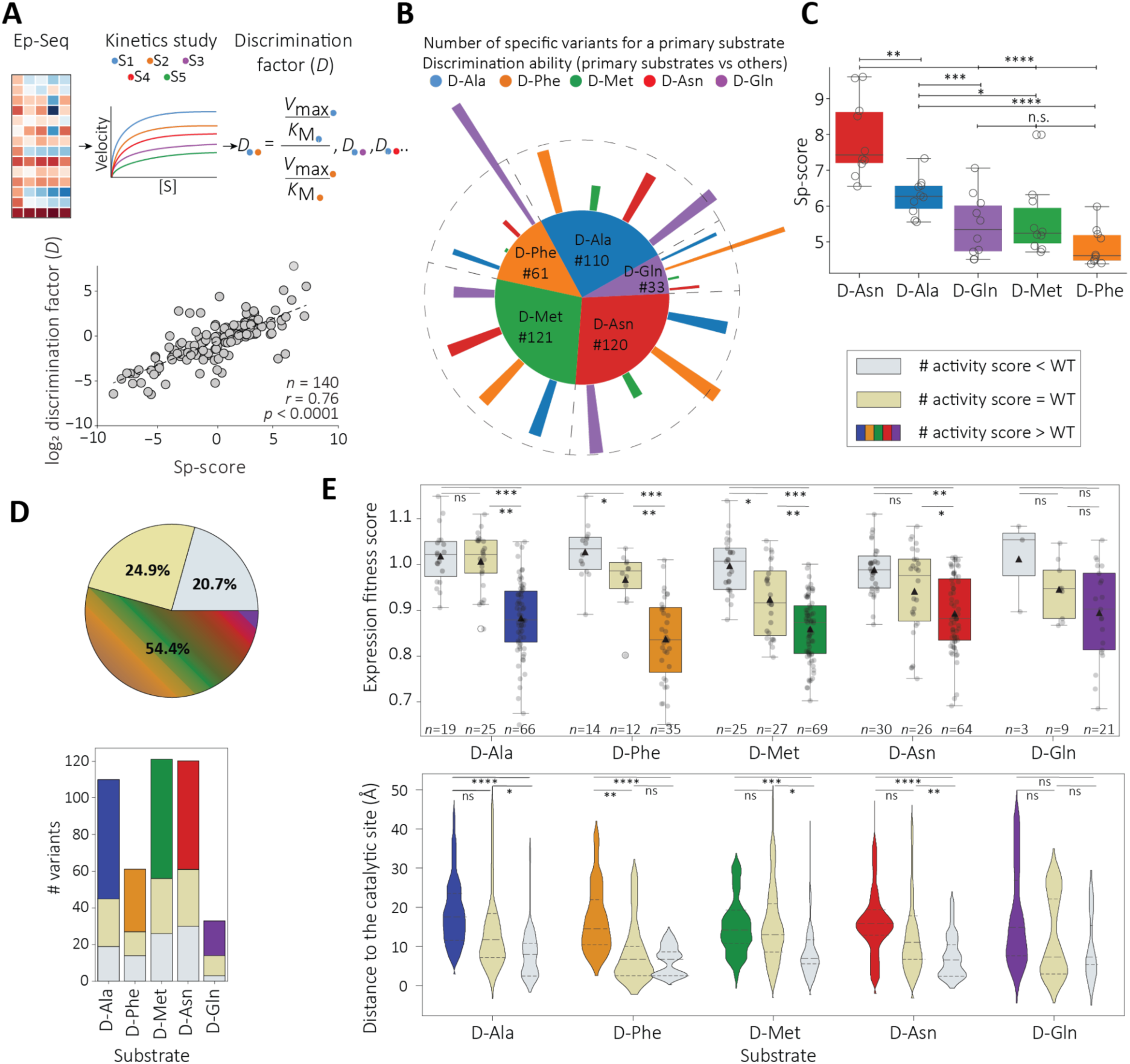
Validation and global analysis of substrate specificity scores. (**A**) Comparison of EP-Seq-derived specificity scores (Sp-scores) with single-clone discrimination factors (D-factors) determined by Michaelis-Menten kinetic analysis across 140 variant-substrate pairs. Dashed line: linear regression; *r*: Pearson correlation coefficients, n: number of datapoints, *p*: p-values two-tailed. (**B**) Pie chart showing the number of specificity-conferring unique mutations identified for each substrate. Outer bar heights represent the intensity of discrimination of variants specific for the substrate shown in the pie chart against all non-cognate substrates. (**C**) Box plot showing Sp-score distributions for the 10 most selective variants per substrate. (**D**) Pie chart (top) summarizing different functional categories of substrate-specific variants: those with reduced (grey), unchanged (beige) or increased (multi-colored) catalytic activity toward their cognate substrate. The bar plot shows the distribution of these categories across each of the five substrate datasets. (**E**) (Top) Box plots of expression fitness scores (proxy for stability) for each substrate-specific variant set grouped by catalytic activity relative to WT. (Bottom) Violin plots showing the distribution of distances from each mutation site to the catalytic center grouped by substrate and catalytic activity relative to WT. n.s. not significant. * indicates *p* < 0.05, ** indicates *p* < 0.01, *** indicates *p* < 0.001 and **** indicates *p* < 0.0001.

We next analyzed the distribution of substrate-specific mutations across all five assays. Among 445 unique single-mutant variants with |Sp-score| ≥ 3 towards at least one of the tested substrates, the majority favored D-Met (n = 121), D-Asn (n = 120), or D-Ala (*n* = 110) (**Fig. 4B, pie chart**). The ten most selective variants for the smallest two substrates D-Ala or D-Asn (irrespective of the non-cognate substrate) exhibited significantly higher average Sp-scores (D-Asn: 7.91 ± 1.10, *n* = 10; D-Ala: 6.28 ± 0.54, *n* = 10) as compared to the top ten most specific variants for the three larger substrates, including hydrophobic D-Phe (4.87 ± 0.53, n=10) and D-Met (5.59 ± 1.00, n = 10), or the polar D-Gln (5.45 ± 0.88, n = 10) (**Fig. 4C**). These trends suggest that the strongest substrate-selective effects arise when mutations act to exclude sterically larger non-cognate substrates.

To quantify patterns of substrate discrimination, we next calculated the mean Sp-score values for each substrate pair by multiplying the number of selective variants (|Sp-score| ≥ 3) for a given substrate pair by their average Sp-score, and normalizing by the total number of variants selective for the same primary substrate. This produced a single value reflecting both the frequency and intensity of discrimination across all substrate pairs (**Fig. 4B, bars**). Discrimination was most pronounced when substrate pairs differed substantially in terms of size or polarity. For example, D-Ala– and D-Asn-specific variants most effectively excluded the bulkier substrates D-Phe and D-Gln. Conversely, D-Phe-specific variants discriminated strongly against D-Gln but less so against the smaller substrates D-Ala, D-Asn, and D-Met. A similar pattern held for D-Gln-specific variants, which discriminated against D-Phe but not against smaller substrates. These trends suggest that accommodating the larger substrates requires an open active site that also admits smaller substrates, limiting selectivity based on steric exclusion. The strong mutual discrimination between D-Phe and D-Gln likely reflects changes in electrostatic interactions. Finally, D-Met-specific variants exhibit moderate and balanced selectivity across all four other substrates, consistent with its intermediate size and chemical features (**Fig. 4B, bars**).

A detailed analysis of the amino acid substitutions in the most selective variants revealed distinct trends, especially for D-Ala and D-Asn. Among the top 10 variants selective for D-Ala, 7 introduced residues bulkier than the WT residue, typically tyrosine (Y) or tryptophan (W) at positions near the catalytic site (average distance = 4.62 Å ± 2.54; **Table S4**). This suggests that steric exclusion of larger non-cognate substrates is a dominant mechanism for enhancing selectivity on D-Ala. In contrast, only 2 of the top 10 D-Asn-selective variants showed increased bulkiness while 8 introduced charged or polar side chains (**Table S4**), suggesting that substrate exclusion alone was insufficient to generate selectivity on D-Asn, which the wild-type enzyme poorly accommodates (*K*_M_ > 10 mM).

Analysis of the 10 most selective variants on D-Phe, D-Met, and D-Gln, which exhibited lower average Sp-scores than the 10 most selective variants on D-Ala or D-Asn, did not reveal any consistent substitution trends (**Table S4**), suggesting that specificity for these substrates arises through more complex or context-dependent mechanisms.

Together, these results indicate that steric clashes and electrostatic interactions play dominant roles in driving substrate specificity. This observation supports the broader idea that positive selection (i.e., enhancing activity on a cognate substrate) is typically a slow and incremental evolutionary process. By contrast, specificity can emerge more rapidly through negative selection, where mutations exclude non-cognate substrates via structural or electrostatic incompatibility^3,4,26,27^. This strategy also applies in many laboratory evolution campaigns, where undesired activity is penalized while variants enhancing specificity are often reached by excluding secondary substrates from the catalytic site^28–30^.

### Evolutionary interplay between substrate specificity, catalytic activity and structural stability

Engineering enzymes for improved activity often compromises structural stability, a well-established trade-off reflecting the delicate balance between function and fold^17,31–34^. To investigate how this balance shifts during evolution of substrate specificity, we analyzed substrate-specific DAOx variants by combining the catalytic activity data with yeast display expression scores, known to be a valid proxy for folding and stability^17,35–37^.

Among the 445 specificity-conferring single mutants, 24.9% (n = 111) retained catalytic activity toward the primary substrate comparable to WT. This suggests that specificity can emerge through functionally neutral drift. These mutations were mildly destabilizing (average expression score: 0.95 ± 0.08; WT score: 1) and generally located away from the active site (average distance: 13.60 ± 10.29 Å) (**Fig. 4 D-E, beige**).

An additional 20.7% (n = 92) of substrate-specific variants exhibited reduced catalytic activity toward the primary substrate, but were similarly stable or slightly more stable than WT (average expression score: 1.00 ± 0.06; WT score: 1). These mutations tended to occur closer to the active site (average distance: 7.96 ± 6.61 Å) and reflect a trade-off in which enhanced specificity is achieved at the cost of catalytic efficiency (**Fig. 4 D-E, gray**).

The remaining 54.4% of variants (n = 242) increased both specificity and catalytic activity but imposed the most substantial reduction in stability, with the lowest expression scores (0.87 ± 0.08) and the greatest average distance from the active site (average distance: 17.1 ± 8.81 Å) **(Fig. 4 D-E, colored**). These results highlight how functional gains can often originate from distal mutations, even when accompanied by destabilizing effects. Notably, the most impactful specificity-conferring mutations clustered near the active site (**Fig. 3C**), suggesting that large shifts in substrate preference often involve functional compromises, passing through regions of reduced fitness along the evolutionary trajectory.

### Single mutations reshape substrate preference via distinct structural mechanisms

To understand in more detail the structural mechanisms underlying substrate specificity, we characterized the catalytic properties of 30 single-mutant DAOx variants chosen for their high Sp-scores. Each variant, along with WT-DAOx was expressed on yeast and assayed for Michaelis-Menten kinetic parameters against all five substrates. We determined *V*_max_ and *K*_M_ values and compared them to WT values to assess changes in substrate affinity and catalytic rate (**Table S5**).

Among D-Ala-selective variants, F58W and F58Y exhibited the highest specificity. Located at the entrance of the substrate tunnel, these mutations increase steric hindrance, restricting access to larger substrates. While WT-DAOx already contains a bulky phenylalanine at this position, introducing even larger residues (Trp or Tyr) significantly shifted the enzyme’s preference toward D-Ala. In the F58W variant, the D-factor for D-Ala over D-Phe increased by >200-fold (D_D-Ala, D-Phe_: F58W = 42.9, WT = 0.2), while an average ∼20-fold increase in D-factor was observed in F58W for D-Ala over D-Met and D-Asn (**Fig. 5 i, Table S5**).

**Figure 5.**
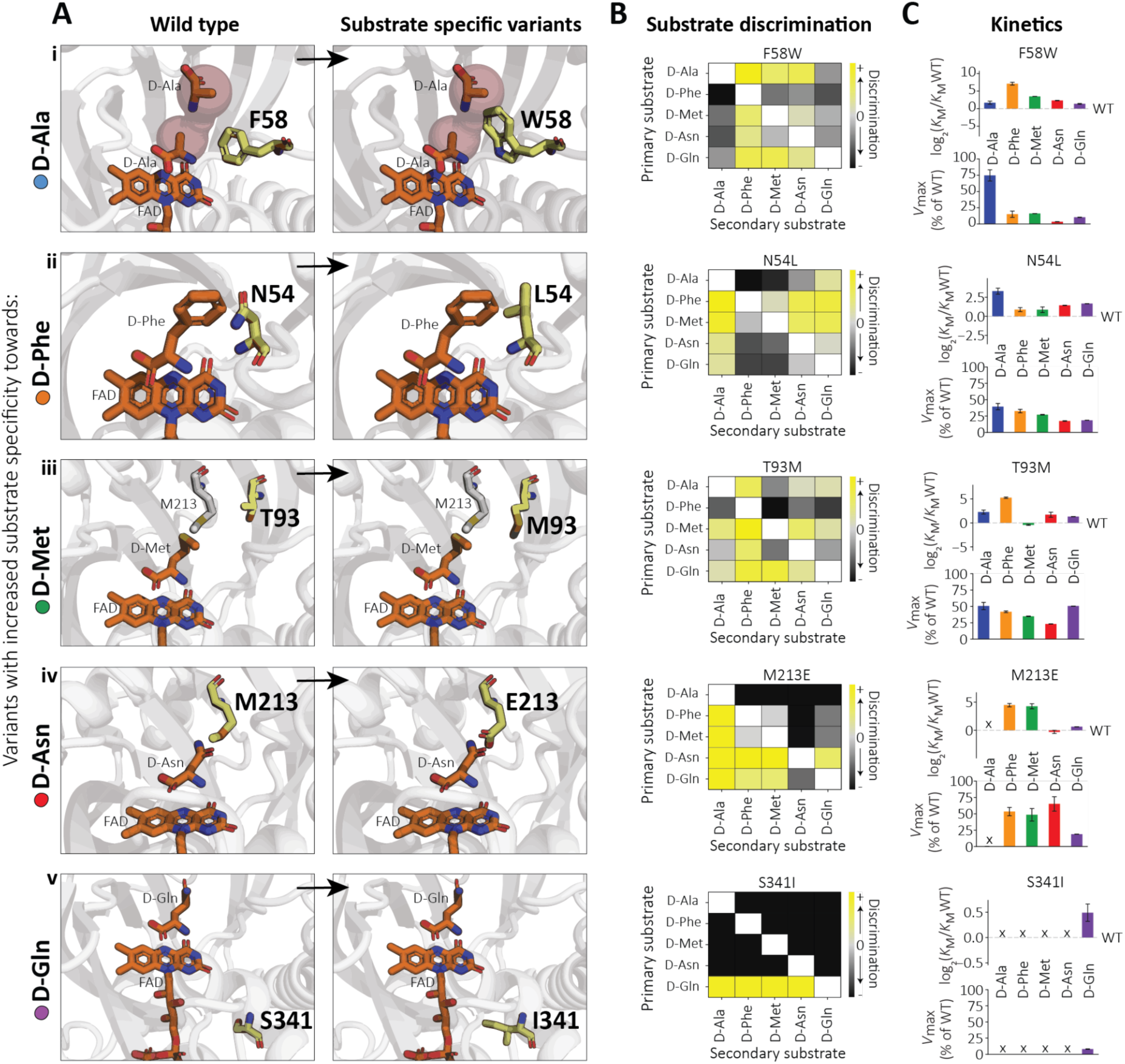
Structural and kinetic insights into substrate-specific DAOx variants. (**A**) Top substrate-selective mutations for each of the five tested substrates mapped onto the three-dimensional structure of DAOx (PDB ID: 1C0P). The FAD cofactor and D-Ala are shown as orange sticks; oxygen, nitrogen, and sulfur atoms are colored red, blue, and yellow, respectively. When relevant, the substrate access tunnel is depicted as light red spheres^40^. Wild-type or substituted residues at the mutation sites are shown in yellow sticks; all other residues are shown in gray. (**B**) Heatmap of substrate discrimination values between all pairwise combinations of primary (y-axis) and secondary (x-axis) substrates, quantifying the degree of specificity conferred by each mutation. (**C**) Catalytic profiles of selected mutants expressed on the surface of the yeast. Relative changes in *K*_M_ and *V*_max_ values compared to the wild-type enzyme for each of the five substrates, illustrating how specificity is tuned through alterations in substrate affinity and catalytic turnover. X indicates no measurable activity, with signal below the detection limit of the assay.

For the D-Phe substrate, N54L emerged as the most selective variant. In the WT-DAOx catalytic site, Asn54 coordinates a water molecule^25^ and orients substrates through its polar side chain that projects toward the center of the catalytic pocket. Replacing Asn54 with hydrophobic Leu without increasing steric bulk likely alters local solvent interactions, enhancing binding of large hydrophobic substrates. This enhances compatibility with D-Phe and D-Met over the smaller D-Ala (D_D-Phe, D-Ala_: N54L = 22.9, WT = 5.1). The structurally similar N54I showed a comparable selectivity profile as N54L (D_D-Phe, D-Ala_: N54I = 23.3, WT = 5.1) (**Fig. 5 ii, Table S5**).

On D-Met, the T93M mutation located just outside the active site increased selectivity by modestly improving D-Met affinity while broadly reducing catalytic efficiency on the other substrates. A similar effect was observed in T56M, likely due to stabilization of nearby contacts involving Met213. For T93M, discrimination between D-Met and D-Phe increased ∼40-fold (D_D-Met, D-Phe_: T93M = 39.8, WT-DAOx = 0.9,) (**Fig. 5 iii, Table S5**).

For the polar substrate D-Asn, the charged substitution M213E conferred strong selectivity, consistent with prior reports implicating this position in electrostatic tuning of the active site^38,39^. M213E reduced overall enzyme activity but preserved or enhanced affinity for D-Asn and D-Gln, with absolute selectivity against D-Ala and an average ∼30-fold increase in D-factor for D-Asn over the other two hydrophobic substrates D-Phe and D-Met (D_D-Asn, D-Phe_: M213E = 0.7, WT-DAOx = 0.02; D_D-Asn, D-Met_: M213E = 0.73, WT-DAOx = 0.02). The similar variant M213D further reinforced these observations (**Fig. 5 iv, Table S5**).

For the polar substrate D-Gln, we found that the variant S341I exhibited striking selectivity despite the mutation being >8 Å away from the catalytic site and not directly involved in substrate coordination. While *K*_M_ for D-Gln slightly increased, *V*_max_ dropped to ∼10% of WT. No measurable activity was observed for any of the other four substrates, even under extended assay conditions, making this variant fully selective for D-Gln, which is notably one the least favored substrates of WT-DAOx. This highlights how even distal mutations can fine-tune specificity through subtle geometric or allosteric effects (**Fig. 5 v, Table S5**).

To reinforce the validity of our observations, we selected the five DAOx variants that exhibited the most pronounced shifts in substrate specificity toward each of the five tested substrates (F58W, N54L, T93M, M213E, S341I) for soluble expression in *E. coli* and subsequent purification. These variants, together with soluble WT-DAOx, were comprehensively characterized for their Michaelis-Menten kinetics against each of the five substrates used in this study (**Table 1, Fig. S8**). We then calculated discrimination factors and compared them to the corresponding profiles obtained from the yeast-displayed formats (**Fig. S9A-B**). For all unique substrate pairs across these five variants, the determined D-factors showed strong agreement between the soluble and yeast-displayed forms (n = 36, *r* = 0.87, *p* < 0.0001; **Fig. S9B**). All variants retained the specificity profiles observed in the yeast-displayed format. Notably, S341I maintained absolute selectivity for D-Gln, even when tested at concentrations 20-fold higher than WT and under extended incubation times.

**Table 1.** Kinetic parameters of soluble WT– and substrate-specific-DAOx variants tested with each of the five substrates used in this study. Values in parentheses indicate *K*_M_ and *k*_cat_ as a percentage of the corresponding value for WT-DAOx. X indicates no measurable activity, with signal below the detection limit of the assay.

| DAOx soluble variants | D-Alanine |  | D-Phenylalanine |  | D-Methionine |  | D-Asparagine |  | D-Glutamine |  |
| --- | --- | --- | --- | --- | --- | --- | --- | --- | --- | --- |
| | $K_M$ [mM]<br>(%WT) | $k_{cat}$ s<sup>-1</sup><br>(%WT) | $K_M$ [mM]<br>(%WT) | $k_{cat}$ s<sup>-1</sup><br>(%WT) | $K_M$ [mM]<br>(%WT) | $k_{cat}$ s<sup>-1</sup><br>(%WT) | $K_M$ [mM]<br>(%WT) | $k_{cat}$ s<sup>-1</sup><br>(%WT) | $K_M$ [mM]<br>(%WT) | $k_{cat}$ s<sup>-1</sup><br>(%WT) |
| WT | 5.62 ±0.81<br>(100) | 1.47±0.06<br>(100) | 2.56±0.35<br>(100) | 1.93±0.05<br>(100) | 2.02±0.16<br>(100) | 1.37±0.05<br>(100) | 47.06±0.81<br>(100) | 0.84±0.11<br>(100) | 18.84±1.99<br>(100) | 0.81±0.05<br>(100) |
| F58W | 10.43±1.33<br>(190) | 2.21±0.20<br>(150) | 43.73±7.55<br>(1750) | 0.22±0.03<br>(11) | 8.90±0.72<br>(445) | 0.43±0.00<br>(31) | 47.90±9.34<br>(102) | 0.06±0.01<br>(7) | 30.70±1.57<br>(165) | 0.07±0.00<br>(9) |
| N54L | 22.04±2.52<br>(400) | 1.20±0.17<br>(82) | 1.60±0.04<br>(63) | 1.07±0.02<br>(56) | 1.35±0.12<br>(68) | 0.90±0.02<br>(65) | 59.29±8.65<br>(126) | 0.32±0.01<br>(39) | 36.75±1.61<br>(197) | 0.21±0.02<br>(26) |
| T93M | 16.94±2.41<br>(302) | 0.15±0.03<br>(10) | 13.75±0.26<br>(543) | 0.08±0.00<br>(4) | 0.74±0.01<br>(37) | 0.12±0.00<br>(8) | 41.02±14.05<br>(87) | 0.03±0.01<br>(4) | 20.86±1.10<br>(111) | 0.06±0.00<br>(7) |
| M213E | 29.71±15.59<br>(518) | 0.13±0.06<br>(9) | 21.52±1.04<br>(847) | 1.85±0.09<br>(96) | 9.99±1.3<br>(499) | 1.75±0.06<br>(127) | 17.26±0.98<br>(37) | 1.30±0.09<br>(156) | 13.09±0.86<br>(70) | 0.22±0.01<br>(27) |
| S341I | X | X | X | X | X | X | X | X | 9.08±0.25<br>(51) | 0.02±0.00<br>(2) |

### Distal mutations as modulators of substrate specificity

Distal mutations can significantly influence enzyme activity and substrate specificity, although their identification and mechanistic roles are challenging to detect. These effects are often mediated through conformational or allosteric changes and are highly context-dependent^41–43^. Previous deep mutational scanning studies have reported contrasting patterns. In some cases, specificity-determining mutations are mainly distal (>9 Å from the catalytic site)^16^, while in others, they cluster within the first shell residues around the active site^13,15,44^.

In DAOx, we found that the strongest specificity-shifting mutations were predominantly located within 8 Å of the catalytic site (**Fig. 3C**), yet these accounted for only 30% (n = 101) of all such mutations. The remaining 70% (n = 231) were scattered across more distal regions of the enzyme. Notably, we identified a hotspot along α-helix 10 (residues 248–263), where positions 254, 258, and 262 were among the most frequently mutated positions affecting specificity (**Fig. 6A**). These residues occupy alternating helical turns and project their side chains in the same direction, forming interactions with β-strands 9–11 (residues 212–235). This region shapes the catalytic cleft and contains critical substrate-coordinating residues, including M213, S215, and P221 (**Fig. 6B**) which are among the most important positions for substrate specificity across all tested substrates (**Fig. 3D, Fig. S7**).

**Figure 6.**
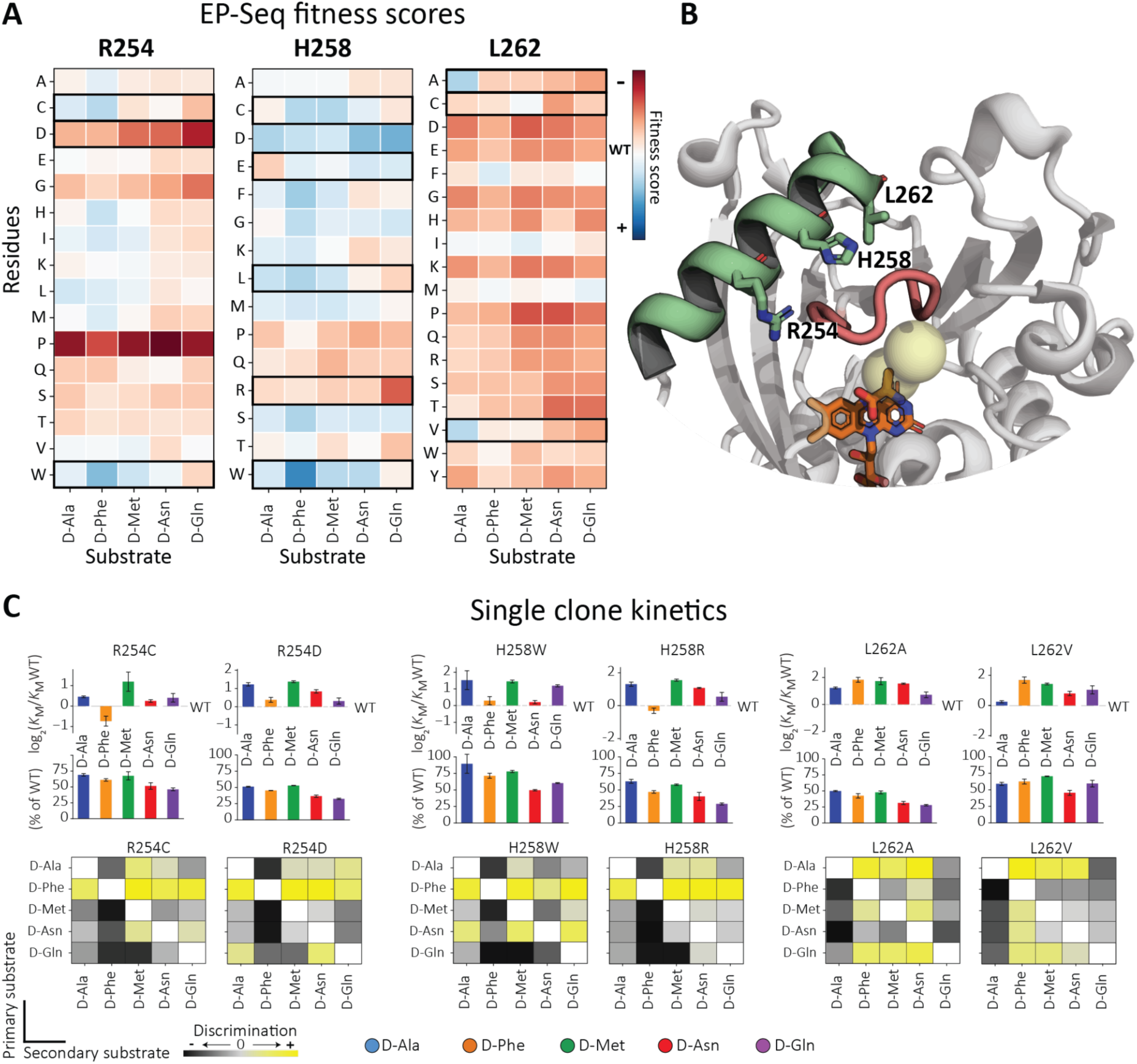
α-Helix 10 as an allosteric site modulating substrate specificity in DAOx. (**A**) Heatmaps of EP-Seq fitness scores for single-point mutants at positions 254, 258, and 262 across all five substrates (D-Ala, D-Phe, D-Met, D-Asn, and D-Gln), highlighting position-specific effects on substrate preference. (**B**) Structural visualization of positions 254, 258, and 262 on α-helix 10, showing side chains oriented toward β-strands 9–11 (residues 212– 235). The loop between β-strands 10 and 11 is shown in red. The FAD cofactor and D-Ala are rendered as sticks (oxygen in red, nitrogen in blue). The substrate tunnel is indicated by light yellow spheres. This view illustrates how distal mutations can alter catalytic site geometry through long-range interactions. (**C**) Kinetic analysis of six representative yeast displayed variants (two per position), showing changes in reaction velocity and substrate affinity across the five tested substrates.

For these distal positions, we characterized two variants per site at the single clone level. At positions 254 and 258, substitutions such as R254C/D and H258W/R resulted in a twofold increase in substrate selectivity toward the bulky hydrophobic substrate D-Phe over all other tested substrates. In contrast, substitutions at position L262 (L262V/A), which introduce smaller hydrophobic side chains, led to a twofold increase in specificity for the smaller substrate D-Ala over all other substrates, with the exception of D-Gln. These results show how distal mutations outside the catalytic site can reshape the protein’s structural network, subtly modifying the catalytic pocket’s geometry and influencing substrate preference. The close proximity of α-helix 10 to the dimer interface hints that oligomerization-induced structural reorganization can further modulate the catalytic site and contribute to substrate specificity.

### Combinatorial mutations enable control over substrate specificity

To further refine substrate specificity for engineering applications, we tested whether combining substrate-specific mutations would produce additive or synergistic effects on substrate preference (**Fig. 7**). Since the most selective single-point mutations were clustered within the catalytic site and also enhanced structural stability, we hypothesized that combining these mutations could further sharpen substrate specificity without significantly compromising stability or expression levels.

**Figure 7.**
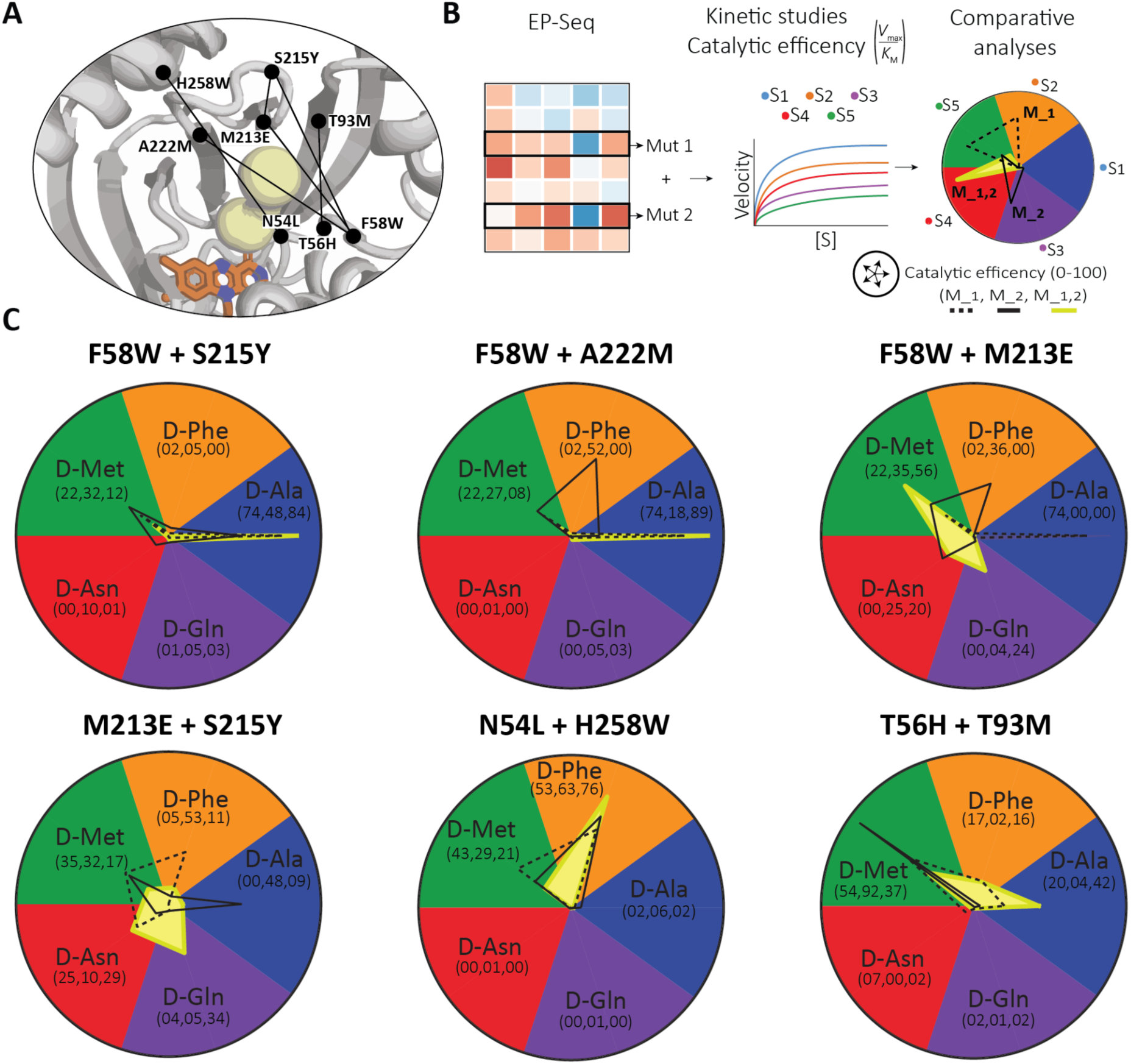
Rational engineering of substrate specificity through combinatorial mutations. (**A**) Structural mapping of single-mutant sites selected for double mutant design in DAOx (PDB: 1C0P). Pairs were chosen based on complementary substrate specificity profiles. (**B**) Workflow illustrating the rational design process: substrate-specific single mutants identified by EP-Seq were first characterized individually and then combined into double mutants. The catalytic efficiency (V_max_/*K*_M_) of each double mutant was measured across all five substrates and compared to the corresponding single mutants to assess additive or synergistic effects on substrate preference. (**C**) Radar plots summarizing the catalytic efficiency profiles of single and double mutants. For each substrate, the percentage of catalytic efficiency is indicated in parentheses. Curves show the performance of single mutant 1 (dashed black line), single mutant 2 (solid black line), and the double mutant (solid yellow filled shapes), highlighting shifts in substrate specificity.

We constructed six double mutants and measured their kinetic parameters across all five substrates, comparing the specificity profiles to those of the corresponding single mutants (**Fig. 7A-B**). For example, the F58W mutation that introduced strong steric hindrance at the substrate tunnel and consistently reduced activity toward D-Phe. When combined with mutations that favor D-Ala such as S215Y or A222M, we observed a sharp drop-in activity for all other substrates and a pronounced enhancement in D-Ala specificity. By contrast, combining F58W with M213E, which introduces a negative charge at the active site, abolished D-Ala activity and shifted specificity toward the polar substrates D-Asn and D-Gln (**Fig. 7C, top**). The M213E/S215Y double mutant further intensified this specificity shift, redirecting enzyme activity almost entirely toward D-Asn and D-Gln, two substrates that are only minimally processed by WT-DAOx.

We also combined N54L and H258W, both of which favor D-Phe individually, and observed a strong additive effect: the double mutant retained over 75% catalytic efficiency for D-Phe while showing greatly reduced activity on all other substrates. However, not all combinations were additive. The T56H/T93M mutant, designed to enhance specificity on D-Met, maintained activity on multiple hydrophobic substrates without a clear preference, demonstrating that certain mutations can interfere with each other’s effects (**Fig. 7C, bottom**).

## Conclusions

Substrate specificity lies at the intersection of an enzyme’s chemical and biophysical characteristics. Once viewed as a static feature dictated solely by rigid active-site geometry, substrate specificity is now understood as an emergent property arising from conformational flexibility and long-range intramolecular contact networks. Enzymes are highly dynamic and sample multiple conformations governed by multiple weak interactions that collectively define substrate recognition ^42,43,45,46^. Despite these insights, encoding this complexity through enzyme design remains a major challenge due in large part to our limited understanding of how distant mutations exchange information and influence the catalytic geometry.

Some work has successfully enhanced specificity by designing rigid substrate-binding sites at the cost of overall activity, and later recovering activity back through directed evolution^47–49^. But concrete principles derived from experimental data describing how specificity is achieved during evolution remain elusive. Here, we contribute to this understanding by presenting a comprehensive, single-mutation resolution map of substrate specificity in the D-amino acid oxidase from *Rhodotorula gracilis*. By leveraging our high-throughput EP-Seq method and exploiting the enzyme’s natural promiscuity in accommodating chemically diverse substrates ^38,50^, we systematically mapped how the mutational fitness landscape shifts across substrates with distinct properties, offering new insights into evolutionary routes to enzyme specificity.

Our results show that mutations near the catalytic site contribute most significantly to substrate specificity. However, these high-impact mutations often come at a substantial cost to catalytic activity. This supports a key evolutionary principle: natural enzymes can rapidly evolve sharp specificity via negative selection mechanisms (steric hindrance or electrostatic repulsion) as long as the functional cost is tolerable. However, other evolutionary routes toward specificity are also available. We identified hundreds of distal mutations that, although generally weaker in magnitude, tend to preserve catalytic activity better and allow specificity to evolve through neutral or slightly biased drift.

Beyond these evolutionary insights, our work also establishes EP-Seq as a robust platform for high-throughput enzyme engineering. The DMS-based Sp-score metric presented here shows a strong correlation with discrimination factors calculated from single-clone kinetic measurements, for both yeast-displayed and soluble variants. Among the strongest single mutations, variants such as F58W, and M213D/E shifted substrate preference by >200-fold. One variant (S341I) achieved exclusive selectivity for D-Gln, one of the WT enzyme’s least favored substrates. We further show that combining mutations with complementary specificity effects can further refine substrate discrimination, offering a powerful strategy for engineering highly selective biocatalysts.

## Acknowledgements

This work was supported by the University of Basel, ETH Zurich, the SNF-NCCR in Molecular Systems Engineering, and an SNSF Grant (200021_191962 and 10004516) to M.A.N. Figure 1 and supplementary figure 9 were partially created using Biorender.com.

## Contributions

R.V. and M.A.N. conceived the study and drafted the manuscript. R.V. carried out the practical work and computational analyses. S.B. optimised and performed the expression and purification of the soluble DAOx variants. C.K. contributed to the conceptualization and optimization of the workflow. M.A.N. secured funding and administered the project.

## Competing interests

The authors declare no competing interests.

## Notes

https://doi.org/10.5281/zenodo.15846928

### Competing Interest Statement

The authors have declared no competing interest.

