## Supplementary Information for "Decoding Substrate Specificity in a Promiscuous Biocatalyst by Enzyme Proximity Sequencing"

#### Materials and Methods

##### Expression and Surface Display of DAOx Monoclonal Populations and Variant Library

The pYDKan_WT-DAOx plasmid and DAOx variant library used in this study were previously constructed and described[1]. The plasmids were transformed into *Saccharomyces cerevisiae* strain EBY100 using the lithium acetate method[2], and positive colonies were selected on synthetic defined (SD) plates or liquid medium containing 2% (w/v) glucose and lacking tryptophan (-Trp). As previously described[1], to induce expression and surface display of the Aga2-DAOx fusion proteins, positive transformants were first cultivated in SD -Trp liquid medium with 2% (w/v) glucose at 30 °C for 24 h with shaking at 200 rpm, until reaching an $OD_{600}$ of approximately 8. Cells were then transferred to fresh SD -Trp induction medium containing 0.2% (w/v) glucose, 1.8% (w/v) galactose, and 100 mM citrate/phosphate buffer (pH 7.0) and incubated at 20 °C for 48 h. Following induction, yeast cells were pelleted, washed with PBS containing 0.1% (w/v) bovine serum albumin (BSA), and labeled with antibodies targeting the C-terminal 6×His tag to assess surface display of the fusion proteins.

##### Yeast antibody staining to detect surface displayed DAOx

As outlined in our previous work[1], following induction of protein expression and surface display, yeast cells were washed twice with PBS supplemented with 0.1% (w/v) BSA. Cells were then resuspended at a density of $2 \times 10^6$ cells per 100 µl in the same buffer containing a 1:500 dilution of mouse monoclonal anti-6xHis antibodies (Thermo Fisher Scientific, Cat# MA1-21315, RRID: AB_557403). Samples were incubated with the primary antibody for 30 minutes at room temperature, followed by two additional washes with PBS/BSA. Next, cells were resuspended in PBS/BSA(0.1%) at the same concentration ($2 \times 10^6$ cells/100 µl) and incubated with a 1:500 dilution of Alexa Fluor™ 594-conjugated goat anti-mouse secondary antibody (Thermo Fisher Scientific, Cat# A-11005, RRID: AB_2534073) for 30 minutes at 4 °C. After incubation, samples were washed twice more and prepared for analysis. Flow cytometry was performed using an Attune NxT cytometer (Thermo Fisher Scientific) equipped with 488 nm and 561 nm lasers. Surface expression of DAOx was quantified by recording and analyzing 10,000 events per sample.

##### Soluble Expression of WT-DAOx

The wild-type D-amino acid oxidase was expressed in Escherichia coli. The gene encoding the wild-type enzyme was cloned into the pET28 bacterial expression vector. Plasmids were transformed into BL21 (DE3) cells and plated on kanamycin-selective LB plates. From a single colony, a liquid LB Kanamycin culture (3 mL) was started and grown overnight at 37 °C with shaking (200 rpm). One milliliter of this overnight culture was used to inoculate 100 mL LB Kan and allowed to grow for 2.5 hours at 37 °C to an $OD_{600}$ of 0.5. To induce expression, IPTG was added to a final concentration of 0.5 mM, and the culture was grown overnight for 18 hours at 25 °C with shaking (200 rpm). The following day, cells were centrifuged at 4000 rpm for 15 minutes, and the supernatant was discarded. The pellet was placed on ice for 15 minutes and then resuspended in 10 mL of lysis buffer (50 mM TRIS, pH 8.0,

50 mM NaCl, 0.1% (v/v) Triton X-100, 5 mM MgCl2). Cells were lysed by sonication for 10 minutes (Sonifier cell disruptor Branson Digital Sonifier, USA) and then centrifuged at 18,000 g for 20 minutes. The supernatant was loaded onto a gravitational flow column packed with 2 mL of HisPur Ni-NTA resin. Columns were washed with 6 column volumes of PBS buffer first and then 6 column volumes of PBS buffer containing 20 mM imidazole. Finally, the bound protein was eluted in 5 mL PBS buffer containing 500 mM Imidazole. Elution fractions were concentrated using Vivaspin concentration columns (10 kDa cutoff). Subsequently, purity was increased, and residual salts from the affinity purification protocols were removed by size exclusion chromatography at a speed of 0.5 mL/min. The most abundant fraction was collected and the purity and molecular weight of the eluted protein were tested and confirmed by running 2 µg of eluted protein through a 12% polyacrylamide SDS gel.

#### Kinetic Characterization and Comparison of Soluble and Yeast-Displayed WT-DAOx

Kinetics measurements for soluble RgDAOx wild type were performed through Amplex Red activity assay by incubating the purified wild type enzyme with increasing concentrations of each tested substrates together with 5.6 µM HRP and 100 µM Amplex Red in PBS (pH 7.5). For measurement we used the following enzyme and substrate concentration. D-alanine, D-phenylalanine, D-methionine, D-leucine were tested at 0.27 mM, 0.55 mM, 1.1 mM, 2.2 mM, 4.4 mM, 8.8mM, 17.5 mM and 35 mM with a WT-DAOx concentration of 0.058 nM. D-histidine was tested at 0.55 mM, 1.1 mM, 2.2 mM, 4.4 mM, 8.8mM, 17.5 mM and 35 mM with a WT-DAOx concentration of 0.058 nM. D-serine was tested at 1.1 mM, 2.2 mM, 4.4 mM, 8.8mM, 17.5 mM 35 mM, 70 mM and 140 mM with a WT-DAOx concentration of 0.058 nM. D-Threonine was assayed at 1.1 mM, 2.2 mM, 4.4 mM, 8.8mM, 17.5 mM 35 mM, 70 mM and 140 mM with a WT-DAOx concentration of 0.29 nM. D-glutamine and D-asparagine were assayed at 0.4 mM, 0.8 mM, 1.6 mM, 3.2 mM, 6.3 mM 12.5 mM, 25 mM and 50 mM with a WT-DAOx concentration of 0.29 nM. The fluorescence signal was read over time at 590 nm and converted to concentration of $H_2O_2$ using a calibration curve. The linear range of each reaction was used to extrapolate the reaction velocity and finally plotted versus substrate concentration to fit Michaelis Menten model and extract $K_M$. $k_{cat}$ was then calculated by dividing the measured $V_{max}$ for each respective substrate by the concentration of enzyme used in the reaction.

Yeast cell populations positive for the display of DAOx (1’000’000 for D-Threonine D-glutamine and D-asparagine, 200’000 for D-alanine, D-phenylalanine, D-methionine, D-leucine, D-histidine and D-serine) were assayed for D-amino acid oxidase activity by mixing the cells with, 5.6 µM HRP, and 100 µM Amplex Red and increasing concentrations of each substrate in PBS (pH 7.5). D-alanine, D-phenylalanine, D-methionine, D-leucine were tested at 0.27 mM, 0.55 mM, 1.1 mM, 2.2 mM, 4.4 mM, 8.8mM, 17.5 mM and 35 mM. D-histidine was tested at 0.55 mM, 1.1 mM, 2.2 mM, 4.4 mM, 8.8mM, 17.5 mM. D-serine and D-threonine were tested at 1.1 mM, 2.2 mM, 4.4 mM, 8.8mM, 17.5 mM 35 mM, 70 mM and 140 mM. D-glutamine was assayed at 0.4 mM, 0.8 mM, 1.6 mM, 3.2 mM, 6.3 mM 12.5 mM, 25 mM and 50 mM. D-asparagine was assayed at 0.5 mM, 1 mM, 2.2 mM, 4.4 mM, 8.8 mM, 17.5 mM, 35 mM and 70 mM. The fluorescence signal was read over time at 590 nm and converted to concentration of $H_2O_2$ using a calibration curve. The linear range of each reaction was used to extrapolate the reaction velocity and finally plotted versus substrate concentration to fit Michaelis Menten function and extract $K_M$ and $V_{max}$.

#### DAOx single cell tyramide/peroxidase proximity labeling assay

##### General procedure

The activity of yeast-displayed DAOx wild type was evaluated using a tyramide-based single-cell assay. Cells were diluted to a final concentration of 500 cells/µl and incubated with a 1:200 dilution of Alexa Fluor™ 488 Tyramide Reagent (Thermo Fisher Scientific, B40953), 55 µM horseradish peroxidase (HRP; Sigma-Aldrich, 77332), and the appropriate concentration of D-amino acid substrate, adjusted according to the specific substrate and experimental application. Reactions were carried out at 25 °C for a defined period depending on the assay conditions. Following incubation, cells were pelleted by centrifugation at 13,000 × g for 3 min using a benchtop centrifuge. The pellet was washed twice with PBS containing 0.1% (w/v) BSA and 0.05% TWEEN-20, then resuspended in the same buffer for flow cytometric analysis.

##### Time course tyramide assay

Time course analyses of the tyramide assay were conducted under the same conditions as described above, with each substrate concentration set to its respective calculated $K_M$. Specifically, the concentrations were as follows: 7.3 mM D-alanine, 2 mM D-methionine, 1.5 mM D-phenylalanine, 14.4 mM D-glutamine, and 45.8 mM D-asparagine. For reactions utilizing D-alanine as a substrate, incubation times were set at 5, 10, 20, 30, 40, 50, 60, 70, 80, and 90 minutes, respectively. Reactions involving D-phenylalanine and D-methionine were incubated for 1, 5, 10, 20, 30, 40, 50, 60, 70, 80, and 90 minutes, respectively. Lastly, reactions using D-asparagine and D-glutamine as substrates were incubated for 10, 20, 30, 40, 50, 60, 80, 100, 120, and 140 minutes, respectively.

##### DAOx variants library tyramide assay

Catalytic activity of displayed DAOx variant enzymes was assayed through a single-cell tyramide/peroxidase proximity labeling method. After induction of protein expression and cell surface staining of the yeast population as described above, 25M yeast cells were mixed at a concentration of 500 cells/µl with 1/200 dilution of Alexa fluor™ 488 Tyramide Reagent (Thermo Fisher, B40953), 56.8 µM HRP (Sigma-Aldrich, 77332) and one of the five substrates at concentration equal to $K_M$ in 1xPBS and 0.75% (w/v) sodium alginate. The reactions performed with D-alanine, D-phenylalanine and D-methionine were incubated for 60 minutes at 25°C. The reactions performed with D-glutamine and D-asparagine were incubated for 120 minutes at 25°C. Afterwards, the cells were spun down for 3 min at 13000 g in a table top centrifuge. The cell pellet was washed twice with PBS 0.1% (wt/vol) BSA + 0.05% (vol/vol) Tween 20 and then used for flow cytometry and single cell sorting experiments.

#### Activity level sorting

EBY100 yeast cells expressing DAOx enzyme variants were stained to assess surface expression and assayed for activity using the single-cell tyramide assay. Cells were subsequently sorted on a FACSMelody cell sorter (BD Biosciences) equipped with 488 nm and 561 nm lasers and a 100 µm nozzle, following the protocol described in our previous study[1] with minor modifications. Cells were sorted into pre-wetted 5 ml FACS tubes containing 0.5 ml of 2× concentrated SD -Trp medium supplemented with 4% (w/v) glucose and 1% (w/v) BSA. Yeast cells were first gated for singleton events and then the population divided into four bins based on the level of green fluorescent signal.

Bin 1 was designed in order to include 99% of the population of cells negative to display or displaying inactive DAOx variants, using as reference the fluorescence level of the non-displaying yeast populations. The remaining part of the population of cells was equally divided into three yeast sub-populations with increasing fluorescent signal. We performed the sorting of two independently assayed yeast populations sorting each time between 5 M and 12 M total yeast cells therefore covering the entire variability of the library of 200'000 maximum variants at least 25 fold. After each cycle of sorting, yeast cells part of the same gated population were merged into 50 ml falcon tubes and pelleted 10 min at 4000 g in a table top centrifuge. Afterwards, the supernatant was discarded and the cell pellet resuspended in 10 ml -Trp medium with 2% glucose supplemented with 100 µg/ml Pen Strep. After sorting, all the cell populations were grown for 48h at 30°C shaking at 200 rpm before being sampled into aliquots of 50M cells each and stored at -80°C in 25% (v/v) glycerol.

#### Preparation of DNA Libraries for Illumina Sequencing and Data Analysis

As previously outlined[1], approximately 50 million yeast cells per sorted population were used as starting material for the preparation of Illumina sequencing libraries. Frozen cell pellets stored at −80°C were thawed and incubated at room temperature for 5 minutes, then centrifuged at 13,000 × g for 1 minute in a benchtop centrifuge. The supernatant was discarded, and the cell pellet was resuspended in 250 µl of miniprep resuspension solution (GeneJET Plasmid Miniprep Kit, Thermo Fisher) supplemented with 4 µl of Zymolyase (5 U/µl). Samples were incubated at 37°C for 2 hours with shaking at 900 rpm to digest the yeast cell walls. Following enzymatic treatment, plasmid DNA was extracted according to the manufacturer's protocol for the GeneJET Plasmid Miniprep Kit. The purified plasmid DNA was eluted in 15 µl of nuclease-free water. The region containing the 15N unique molecular identifier (UMI) was amplified by PCR using NEBNext Ultra II Q5 Master Mix (New England Biolabs). Each 50 µl PCR reaction contained 25 µl of master mix, 5 µl of forward primer (1 µM), 5 µl of reverse primer (1 µM), and 15 µl of template DNA. Primers were designed to target the region of interest and to be compatible with Nextera indexing library preparation. To enhance sequence diversity at one end of the amplicon, four staggered forward primers were used in separate PCR reactions, and the resulting products were pooled for sequencing. PCR cycling conditions were as follows: initial denaturation at 98°C for 30 seconds; 18 cycles of 98°C for 10 seconds and 72°C for 30 seconds; followed by a final extension at 72°C for 2 minutes. Reactions were held at 4°C until further processing. PCR products were verified by electrophoresis on a 1% (w/v) agarose gel and purified using a standard DNA gel extraction kit. DNA concentration was measured, and samples were further purified using the DNA Clean and Concentrator-5 kit (Zymo Research). Unique Nextera indexing sequences were then added, samples were pooled, and sequencing was performed on an Illumina NovaSeq 6000 platform. Illumina read counts per sample are detailed in **Table S2**. Demultiplexed sequencing reads were processed using a computational pipeline to extract UMI sequences and align them against a look-up table. UMI sequences were mapped using the BBMap alignment algorithm [3] with the reference sequence file (ill_ref.fa)

```
bbmap.sh in = *.fastq ref = ill_ref.fa out =*.sam
```

BBMap-generated sam files were further processed using the C script pib.c (process Illumina barcodes). This step extracted the sequences and associated quality scores for each UMI mapped by BBMap, saving the output in FASTQ format.

```
./pib *.sam> *.fq
```

Each mapped UMI was then aligned against the sequences stored in the look-up table (lut_m.tsv) using the C script rib.c (read Illumina barcodes). Each barcode was assigned one of the following tags based on the alignment and quality criteria: 0 – not found in the look-up table; 1 – found in the look-up table; 2 – read quality below Q20; 3 – UMI length different from 15 nucleotides. We applied a quality filter at Q20, tagging all UMIs associated with reads of lower quality with tag 2.

```
./rib -t lut_m.tsv -q 20 *.fq > *.tsv
```

Finally, UMI sequences with tag 1 were extracted, sorted alphabetically and grouped by identity.

```
grep^1 *.tsv |sort|uniq -c|sed -E ‘s/^*//; s/ /\t/’>t1sct_*.tsv
```

All Illumina read files were processed using the computational workflow described above. The UMI sequence data and counts from each bin (t1sct_Bin#.tsv) were subsequently merged into a single file using the Jupyter notebook ill_tag1_bins. The C programs employed in this analysis were compiled with GCC version 7.4.0 on a 32-bit processor. Raw sequencing reads are accessible via the NCBI Sequence Read Archive under the Bioproject accession number: PRJNA1289092. The scripts, look up table and output data are available at https://doi.org/10.5281/zenodo.15846928

#### Fitness scores calculation

As in our previous work [1], the number of sequencing reads corresponding to each enzyme variant ($r_v$) was converted into the number of sorted cells for that variant ($c_v$) per sorted bin using the following equation (1):

$$\frac{r_{\mathrm{v}}}{r_{\mathrm{tot}}} = \frac{c_{\mathrm{v}}}{c_{\mathrm{tot}}} \qquad \textbf{(1)}$$

Here, $r_{\mathrm{tot}}$ represents the total number of Illumina reads in the bin, and $c_{\mathrm{tot}}$ is the total number of cells sorted into the same bin. The final activity scores for each variant and substrate were then calculated as the expected fluorescent intensity across all four bins in the experiment. We computed a weighted mean ($\beta$) of the cell counts ($c_v$), using the median fluorescence intensity of the yeast population in each bin ($\omega$) as the weighting factor:

$$\beta = \frac{\sum_{i=1}^{bin} \omega_i \cdot c_{\mathrm{v}i}}{\sum_{i=1}^{bin} c_{\mathrm{v}i}} \qquad \textbf{(2)}$$

The fitness score ($F$) per variant were finally calculated as follow:

$$F = \log_2(\frac{\beta_v}{\beta_{wt}}) \qquad \textbf{(3)}$$

where $\beta_v$ represents the weighted mean expression score of the variant enzyme, and $\beta_{wt}$ corresponds to the score of WT-DAOx. To calculate the final consensus fitness score ($F_{fin}$) for each variant, we computed a weighted mean of the individual fitness scores ($F_v$) obtained from each replicate assay. The weighting factor used was the number of cells ($c_v$) associated with the measured fitness in each experiment (**Eq. 4**).

$$F_{fin} = \frac{\sum_{j=1}^{rep} F_{vj} \cdot c_{vj}}{\sum_{j=1}^{rep} c_{vj}} \qquad \textbf{(4)}$$

##### Normalization of activity fitness scores by expression levels

To account for differences in enzyme display (i.e., expression) levels, and before comparative analyses, activity scores were exponentiated (i.e., inverted from $\log_2$ scale) and normalized to expression scores obtained from our previously published dataset [1]. For this procedure, we included only variants with expression levels exceeding the mean expression of nonsense variants plus one standard deviation, to prevent inflated normalized activity scores due to extremely low expression values.

#### Comparative analysis and Sp-score

To identify mutations altering substrate preference, we calculated z-scores for each missense variant across all pairwise substrate combinations using the formula (**Eq. 5**):

$$\text{z-score} = \text{Sp-score} = \frac{X - \mu}{\sigma} \textbf{(5)}$$

where X is the normalized activity fitness difference of the variant measured between two substrates, and μ and σ are the mean and standard deviation of fitness differences across all variants for the same substrate pair. This metric, termed substrate specificity score (Sp-score), captures both the magnitude and direction of substrate preference changes. Using synonymous mutations as controls, we set a threshold off |Sp|≥3 (three standard deviations) to identify substrate-specific mutations, excluding >99% of synonymous variants.

##### Quantification of substrate discrimination across substrate pairs

The intensity of discrimination for variants specific to a primary substrate against all non-cognate substrates presented in Fig. 4B was calculated in two steps. First, for each substrate pair, we computed the proportion of cognate substrate–specific variants (|Sp-score| ≥ 3) relative to the total number of specific variants for the primary substrate. This proportion was then multiplied by the mean Sp-score of these variants toward the primary substrate within that substrate pair. The resulting metric integrates both the prevalence and magnitude of substrate discrimination.

##### DAOx structural properties calculations from 3D structure

Coordinates of all atoms in DAOx residues were extracted from the PDB structure 1C0P. The minimum distance from each residue to the crystallized substrate D-Ala was calculated and used as a metric for proximity to the catalytic site in the analysis.

##### Validation of EP-Seq fitness scores

Eighteen single mutant DAOx variants were selected based on DMS scores spanning negative to positive values across D-Ala, D-Phe, D-Met, D-Asn, and D-Gln. Genes encoding S48C, K109L, K109W, K109Y, D110I, D110M, D110W, L153R, A187E, A187K, G199E, Q200W, S215A, S215R, T237M, Y238W, S268Y and L310P were synthesized and cloned into the pYDKan plasmid in-frame with the Aga2p coding sequence using a BioXP 3250 synthetic biology workstation (Codex DNA) or generated via PCR and KLD cloning to introduce mutations into the wild-type DAOx sequence. All sequences were codon-optimized for expression in *Saccharomyces cerevisiae* and verified by Sanger sequencing. Final plasmids were transformed into *S. cerevisiae* strain EBY100 using the lithium acetate method. Positive colonies were selected on synthetic defined (SD) agar plates (-Trp) with 2% (wt/vol) glucose and cultivated in -Trp liquid medium with 2% (wt/vol) glucose for 24 h at 30 °C to an $OD_{600}$ ~8 with continuous shaking (200 rpm). Expression and display of Aga2-DAOx wild-type and mutant constructs were induced by transferring cells at $OD_{600}$ = 0.4 to fresh -Trp liquid medium containing 0.2% (wt/vol) glucose, 1.8% (wt/vol) galactose, and 100 mM citrate/phosphate buffer (pH 7). Cells were grown for 48 h at 20 °C, then pelleted, washed with PBS containing 0.1% (wt/vol) BSA, and used for antibody labeling to detect the C-terminal 6×His tag of the displayed fusion protein and confirm expression. The cell populations expressing DAOx wild-type and selected variants were then assayed for activity using the Amplex Red method and the five substrates from the EP-Seq screen (D-Ala, D-Phe, D-Met, D-Asn, D-Gln). A total of 250,000 yeast cells were mixed with 5.6 µM HRP and 100 µM Amplex Red in PBS (pH 7.5), and each substrate was added at the same final concentration used in the EP-Seq workflow (7.3 mM D-Ala, 1.5 mM D-Phe, 2 mM D-Met, 46 mM D-Asn, 14 mM D-Gln). Fluorescence was measured at 590 nm every 50–60 s for at least 60 min and at a temperature of 27-30°C. The linear range of the reaction for all enzyme variants and substrates was determined between 0 and 15 min, and the reaction slope was calculated. The single-clone activity score for each variant was obtained by dividing the reaction slope of the variant by that of the wild-type enzyme. Final scores were averaged from three independent measurements per cell population and plotted against the corresponding EP-Seq activity scores.

#### Kinetic study on yeast displayed DAOx substrate specific variants

Based on our EP-Seq comparative analysis, we selected DAOx variants with the highest z-scores and specificity towards each of the tested substrates. Gene-encoding DAOx variants (**Table S5**) were generated via PCR and KLD cloning to introduce mutations into the wild-type DAOx sequence. All sequences were codon-optimized for expression in *Saccharomyces cerevisiae* and verified by Sanger sequencing. Final plasmids were transformed into *S. cerevisiae* strain EBY100 using the lithium acetate method. Positive colonies were selected on synthetic defined (SD) agar plates (-Trp) with 2% (wt/vol) glucose and cultivated in -Trp liquid medium with 2% (wt/vol) glucose for 24 h at 30 °C to an $OD_{600}$ ~8 with continuous shaking (200 rpm). Expression and display of Aga2-DAOx wild-type and mutant constructs were induced by transferring cells at $OD_{600}$ = 0.4 to fresh -Trp liquid medium containing 0.2% (wt/vol) glucose, 1.8% (wt/vol) galactose, and 100 mM citrate/phosphate buffer (pH 7). Cells were grown for 48 h at 20 °C, then pelleted, washed with PBS containing 0.1% (wt/vol) BSA, and used for antibody labeling to detect the C-terminal 6×His tag of the displayed fusion protein. Yeast cells expressing DAOx wild-type and mutant variants were analyzed by flow cytometry.

Yeast populations were first gated for single events, then divided into four gates based on Alexa Fluor™ 594 fluorescence to cover the entire fluorescence range. A total of 10,000 single cells from each population were analyzed. The median fluorescence value and percentage of cells in each gate were recorded. Each population was assayed in duplicate. A weighted mean ($\beta$) of the cell distribution ($\%c$) was calculated, with the median fluorescence of each gate ($\omega$) as the weighting factor (**Eq. 6**).

$$\beta_{\text{sing. clone}} = \frac{\sum_{i=1}^{gate} \omega_i \cdot \%c_i}{\sum_{i=1}^{gate} \%c_i} \qquad \textbf{(6)}$$

Finally, the single-clone expression score ($F_{sing.clone}$) was calculated by dividing the weighted mean expression of each DAOx variant ($\beta_v$) by the weighted mean expression of the wild-type DAOx ($\beta_{wt}$), both assayed using the same procedure (**Eq. 7**).

$$F_{\text{sing. clone}} = \frac{\beta_{\text{v}}}{\beta_{\text{wt}}} \qquad \textbf{(7)}$$

Yeast populations expressing variants were used to determine the kinetic parameters $K_M$ and $V_{max}$of each DAOx mutant towards the five substrates D-Ala, D-Phe, D-Met, D-Asn, D-Gln.  The number of cells used for each kinetic measurement was fixed to 400 '000 and calculated by measuring the OD of each population using as reference value 1 $OD_{600}$ corresponding to 1M cells in 1 ml. By maintaining fixed the number of cells and taking into account the the expression level of each variant enzyme we wanted to allow not only relative comparative analysis towards substrates for the same variant but also compare the performance of the DAOx single mutants to the wild type enzyme.  Each yeast population was assayed for D-amino acid oxidase activity by mixing the cells with 5.6 µM HRP, 100 µM Amplex Red and increasing concentrations of each substrate in PBS (pH 7.5) at room temperature (25-30°C). Specifically,  D-alanine, D-phenylalanine, D-methionine, were tested at 0.3125 mM, 0.625 mM, 1.25 mM, 2.5 mM, 5 mM, 10mM, 20 mM and 40 mM, while D-asparagine and D-glutamine were assayed at 1.5625 mM, 3.125 mM, 6.25 mM, 12.5 mM, 25 mM, 50 mM, 100 mM, 200 mM.  The fluorescence signal was read for at least 20 min at 590 nm.The linear range of each reaction was determined and used to extrapolate the reaction velocity and finally plotted versus substrate concentration to fit Michaelis Menten function and extract $K_M$ and $V_{max}$ . Reaction velocities were determined for at least two independent replicate reactions at six or more substrate concentrations. These data were used to fit the Michaelis–Menten kinetic model. Due to a technical issue affecting one of the replicate reactions, the R254C variant tested with D-Ala was assessed using a single reaction only. Fitting results were considered reliable and accepted if the coefficient of determination ($R^2$) was approximately 0.90 or higher. In cases where enzyme activity was too low to be accurately detected and the data did not meet the criteria for Michaelis–Menten fitting, reactions were repeated using longer incubation times and/or increased enzyme concentrations. If no improvement was observed, kinetic parameters ($K_M$ and $V_{max}$) were classified as not measurable (X).

#### Discrimination factor calculation

The selectivity ($D$) of an enzyme for a primary substrate is defined as the ratio of the $k_{cat}/K_M$ values for the primary substrate over the same ratio for the secondary substrate. This can be expressed as (**Eq. 8**):

$$D = \frac{(k_{\mathrm{cat}}/K_{\mathrm{M}})\mathrm{sub}_1}{(k_{\mathrm{cat}}/K_{\mathrm{M}})\mathrm{sub}_2} \qquad \textbf{(8)}$$

In this manuscript, when yeast surface-displayed enzymes were used to measure catalytic performance, and the absolute enzyme concentration in the reaction could not be determined, we define the discrimination factor (D) between two substrates as the ratio of $V_{max}$ and $K_M$ (**Eq. 9**)

$$D = \frac{(V_{\mathrm{max}}/K_{\mathrm{M}})\mathrm{sub}_1}{(V_{\mathrm{max}}/K_{M})\mathrm{sub}_2} \qquad \textbf{(9)}$$

This relationship allows to calculate reliable discrimination factor values towards different substrates that reflect the catalytic properties of the soluble counterparts. For each variant we first normalize the $V_{max}$ value to the single clone expression score ($F_{sing.\ clone}$) in order to remove the effect of different levels of expression from the detected reaction velocity. Then we calculated the ratio between $V_{max}/K_M$ for all of the 5 substrates. And the discrimination factor (D) for all possible substrate combinations.

#### Soluble expression of WT and substrate specific DAOx variants

DAOx variants (**Table 1**) were generated by site-directed mutagenesis using PCR and KLD cloning to introduce specific mutations into the wild-type DAOx gene, which was previously cloned into the pET28 expression vector. All constructs were verified by Sanger sequencing  and transformed into *E. coli* NiCo21(DE3) for subsequent expression and purification. Cultures were grown in super optimal broth (SOB, 20 g/L tryptone, 5 g/L yeast extract, 2.5 mM KCl, 10 mM $MgCl_2$, $MgSO_4$, 10 mM NaCl, pH 7.0) at 37 °C with shaking until reaching an O.D.600 of 0.6-0.8. The cultures were then induced using 1 mM isopropyl β-d-1-thiogalactopyranoside (IPTG) at 16 °C, with shaking, for 20 h. After harvesting via centrifugation, the cell pellets were stored at -80 °C. When needed, the cell pellets were thawed and resuspended at 4 °C in BugBuster® protein extraction reagent (MilliporeSigma, Burlington, MA, USA) according to the manufacturers protocol. The resulting cell lysate then underwent centrifugation at 30,000 ×g for 30 min and was passed through a 0.45 µM filter. The clarified lysate was then supplemented with 0.1% (w/v) Tween 20 and 10 mM Imidazole, and incubated with Ni-NTA Resin

(HisPur™, Thermo Fisher Scientific, Massachusetts, United States) at 4 °C, with shaking, for 1 h. Ni-NTA Resin was reconstituted by centrifugation 800 ×g for 5 mins and washed three times with 0.1% (w/v) Tween 20 and 10 mM Imidazole supplemented phosphate-buffered saline (PBS, 137 mM NaCl, 2.7 mM KCl, 10 mM $Na_2HPO_4$, and 1.8 mM $KH_2PO_4$, pH 7.4). The Ni-NTA Resin was then transferred to gravity flow columns (Pierce™, Thermo Fischer Scientific, Massachusetts, United States) and washed using an imidazole gradient, 20-250 mM imidazole-supplemented PBS. Protein-containing elution fractions were then concentrated in centrifugal filters (Amicon®, Merck, Darmstadt, Germany) and further processed using size-exclusion chromatography (SEC, Superrose 12 10/300 GL column, Cytiva, Marlborough, United States) into PBS. SEC elution fractions containing the protein of interest were again concentrated using centrifugal filters and stored at -80 °C, until used in experiments. Protein concentrations were measured by spectrophotometry at 280 nm (NanoDrop 1000, 6 Thermo Scientific, DE, USA).

#### Kinetic Study of Soluble WT and Substrate-Specific DAOx Variants

Soluble WT-DAOx and its variants (Table 1) were used to determine the kinetic parameters $K_M$ and $k_{cat}$ toward five substrates: D-Ala, D-Phe, D-Met, D-Asn, and D-Gln. Each enzyme variant was assayed for D-amino acid oxidase activity by mixing the cells with 5.6 µM HRP, 100 µM Amplex Red, and increasing concentrations of each substrate in PBS (pH 7.5) at room temperature (25–30 °C). WT-DAOx was tested at a concentration of 0.25 nM, while the variants were tested at the following concentrations: N54L at 2 nM, F58W at 2 nM, T93M at 5 nM, M213E at 1 nM, and S341I at 5 nM. For each variant,D-Ala, D-Phe and D-Met were tested at concentrations of 0.3125, 0.625, 1.25, 2.5, 5, 10, 20, and 40 mM, while D-Asn and D-Gln e were assayed at 1.5625, 3.125, 6.25, 12.5, 25, 50, 100, and 200 mM. The fluorescence signal was read for at least 20 min at 590 nm. The linear range of each reaction was determined and used to extrapolate the reaction velocity and finally plotted versus substrate concentration to fit Michaelis Menten function and extract $K_M$ and $V_{max}$ . Reaction velocities were determined for at least three independent replicate reactions at six or more substrate concentrations. These data were used to fit the Michaelis–Menten kinetic model. Fitting results were considered reliable and accepted if the coefficient of determination ($R^2$) was approximately 0.90 or higher. In cases where enzyme activity was too low to be accurately detected and the data did not meet the criteria for Michaelis–Menten fitting, reactions were repeated using longer incubation times and/or increased enzyme concentrations. If no improvement was observed, kinetic parameters ($K_M$ and $k_{cat}$) were classified as not measurable (X).

### Supplementary Notes

#### Note S1. Kinetic characterization of WT-RgDAOx yeast displayed enzyme

RgDAOx is active on a wide range of D-amino acid substrates and for each of them the enzyme exhibits a distinct set of Michaelis-Menten parameters ($K_M$ and $k_{cat}$). Extending EP-Seq and tyramide assay to study the changes in RgDAOx's mutability landscape upon reaction with different substrates requires a detailed characterization of the catalytic features of the displayed enzyme. This ultimately aids selecting the optimal reaction conditions tailored to the specific substrates and ensures comparable settings among reactions. In order to investigate the properties of the displayed RgDAOx we first asked if the displayed enzyme retains the same catalytic properties of the soluble counterpart in terms of affinity and catalytic activity towards the different D-amino acids. We compared the Michaelis constant $K_M$ of the yeast surface displayed enzyme towards different substrates to the one of its soluble counterpart expressed and purified from *Escherichia coli* BL21 cells. $K_M$ serves as an ideal kinetic parameter for this comparison because it remains consistent regardless of the enzyme concentration tested in both systems since it is not possible to precisely quantify and therefore compare the absolute concentration of the enzyme when displayed on the yeast's surface. To this purpose, we selected nine different substrates for DAOx, including D-alanine (D-Ala), D-methionine (D-Met), D-phenylalanine (D-Phe), D-glutamine (D-Gln), D-histidine (D-His), D-threonine (D-Thr), D-asparagine (D-Asn), D-serine (D-Ser) and D-leucine (D-Leu). We assayed the activities of both yeast-displayed and soluble WT-DAOx at various substrate concentrations to determine the Michaelis-Menten $K_M$ constant for each substrate. We detected a high degree of concordance between the $K_M$ values calculated for the two enzymes expressed in different systems (soluble WT-DAOx vs. yeast displayed WT-DAOx, $R^2 = 0.93$) (**Supplementary Table 1**; **Fig. S1**). This allowed us to conclude that the *Rg*DAOx enzyme displayed on the surface of the yeast strain EBY100 maintains its native state, functional properties and substrate preferences even if tethered to the cell wall through the fused Aga2 protein.

#### Note S2. Selection of substrates and conditions for tyramide assay in EP-Seq

We selected a pool of five D-amino acid substrates and experimentally determined the reaction conditions for each, ensuring the best compromise between tyramide signal and reaction incubation time at a substrate concentration equal to $K_M$. The five substrates, D-alanine, D-asparagine, D-methionine, D-phenylalanine, and D-glutamine exhibited high accordance in $K_M$ values between the kinetic measurements performed with soluble and yeast-displayed forms of RgDAOx (**Table S1**). These substrates also span a range of reaction speeds (turnover number) suitable for testing the effective performance of the tyramide assay and more importantly are characterised by diverse physical-chemical features in terms of bulkiness, polarity, and chemical composition of their side chains (**Fig. 1, Fig. S2**). To this purpose we selected three hydrophobic substrates with distinctly different side-chain sizes and chemistries: D-alanine (D-Ala), D-phenylalanine (D-Phe), and D-methionine (D-Met). D-Ala has a small, non-polar methyl group, while D-Phe and D-Met have larger hydrophobic groups—a bulky phenyl ring in D-Phe and a thioether group with significant steric bulk in D-Met. Additionally we selected D-asparagine (D-Asn) and D-glutamine (D-Gln), two substrates with polar side chains that differ by a single $CH_2$ to study how polarity and size of the substrate influence activity of the RgDAOx variants. We then proceeded to carry out the tyramide assay, utilising an initial substrate concentration equivalent to $K_M$ for each substrate, employing a mutant library of DAOx variants. This decision ensures an initial substrate concentration that prompts the reaction to operate at half of its maximum velocity, aligning with the steeper segment of the Michaelis-Menten curve. Consequently, even slight modifications to $K_M$ or the initial reaction velocity will alter the intensity of tyramide labeling on cell surfaces, thereby enhancing the sensitivity of the assay. In contrast, the same activity assay conducted at saturating concentrations of D-alanine ($V_{max}$) lacks sensitivity to mutations that subtly alter $K_M$ (but not $V_{max}$) towards the substrate, as they do not significantly affect the reaction velocity and the resulting tyramide signal acquired by the cells during incubation. For the tyramide assay to be conducted within the EP-Seq with the mutant library and the five substrates, we selected an incubation time of 60 minutes for hydrophobic amino acids and 120 minutes for the two polar amino acids. These incubation times, validated by using the WT-DAOx displaying cells, ensured a consistent shift towards a positive tyramide signal in all cases across the population of cells expressing the enzyme, with a median fluorescence of the fluorescent tyramide assay nearing plateau levels. Incubation times exceeding 120 minutes were avoided to prevent cell precipitation and nonspecific tyramide signaling in the negative population (**Fig. S2**).

#### Supplementary Figures

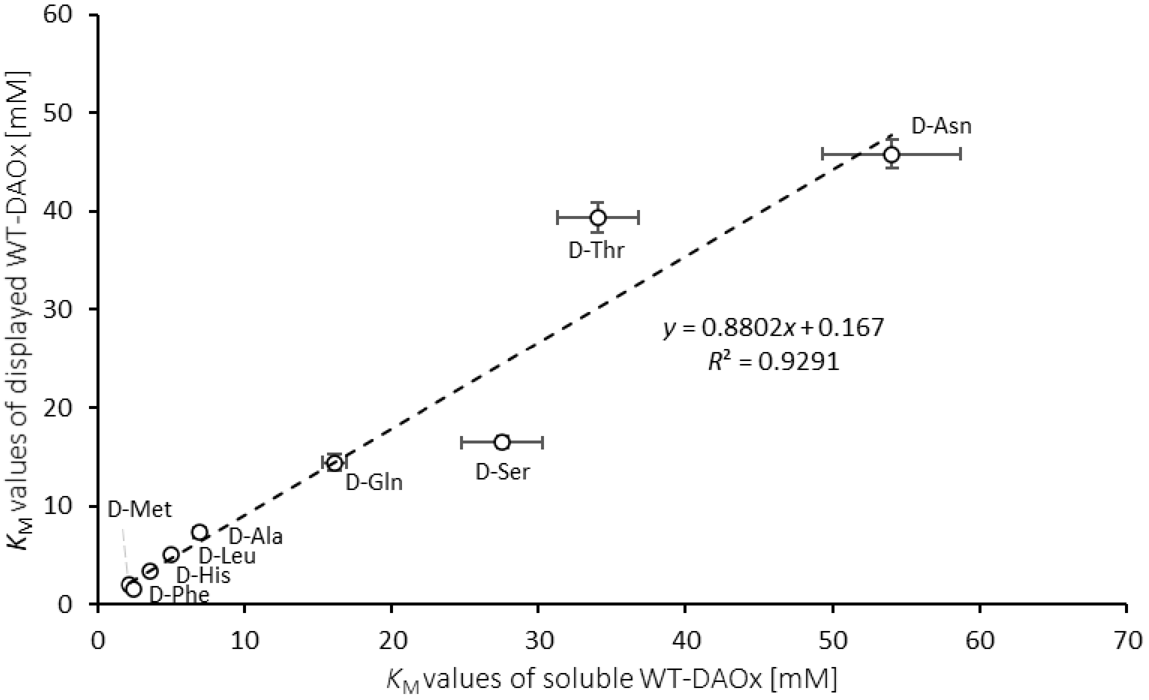

**Figure S1. Comparison of catalytic properties of soluble and yeast-displayed WT-DAOx.**

Linear correlation between the $K_M$ values towards nine different substrates of soluble and yeast displayed wild type DAOx. $K_M$ values for each substrate tested both with displayed and soluble DAOx are reported in Supplementary table 1. Dashed line: linear regression; $R^2$: coefficient of determination.

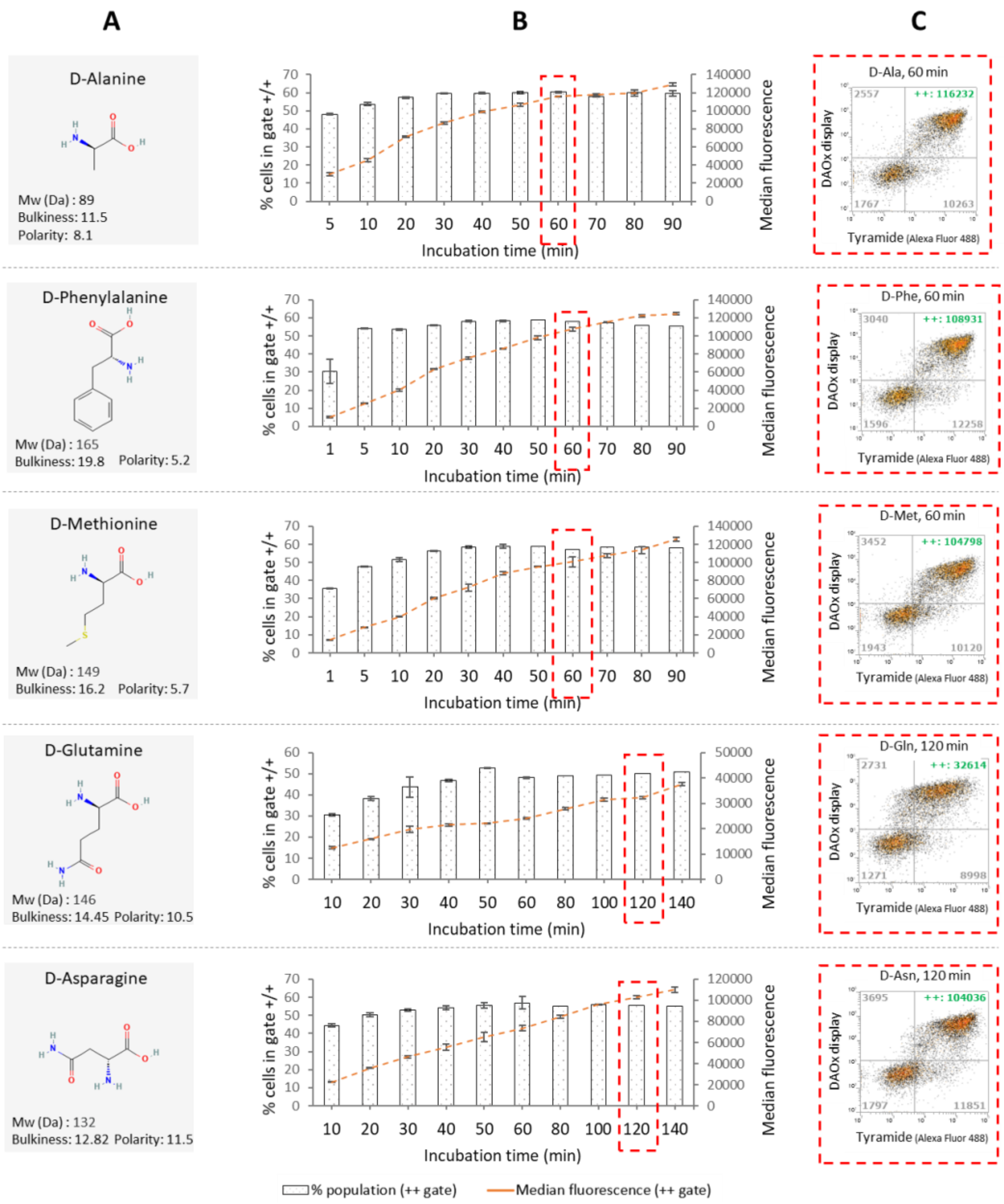

#### Figure S2. Time-course tyramide assay of yeast displayed WT-DAOx

(**A**) Name, chemical structure and information about each of the five DAOx substrates used in the EP-Seq workflow. (**B**) Time course tyramide assay with yeast cells displaying wild type DAOx and five different D-amino acid substrates at an initial concentration equal to $K_M$. For each substrate, a different range of incubation time intervals was chosen based on the calculated $k_{cat}$. D-alanine, D-phenylalanine, and D-methionine (with $k_{cat}$ between 59 and 74 $s^{-1}$) were incubated from 1 to 90 minutes for a total of 9 or 10 time points. D-glutamine and D-asparagine (with $k_{cat}$ of 11 and 18 $s^{-1}$) were incubated for 10 to 140 minutes for a total of 10 time points. Per substrate, the final selected incubation time point used in the deep mutational scanning tyramide screening of the DAOx mutant library is highlighted by the dashed red. The selected time points ensure tyramide signal acquisition close to saturation levels for each amino acid, maximizing the sensitivity of the assay. Each datapoint of the assay was measured from two experimental replicates. (**C**) A raw flow cytometry plot illustrating the population shift along the DAOx display and tyramide signal axis per each substrate and each selected condition is shown.

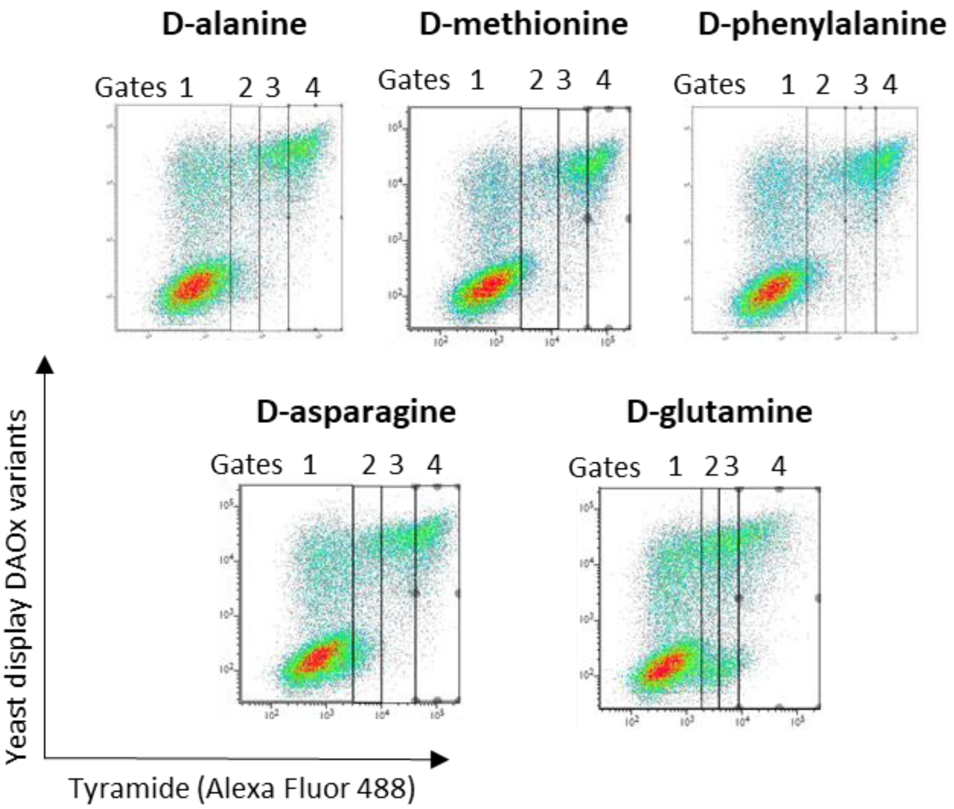

**Figure S3. FACS gating strategy for each of the five EP-Seq tyramide assays performed in the study.**

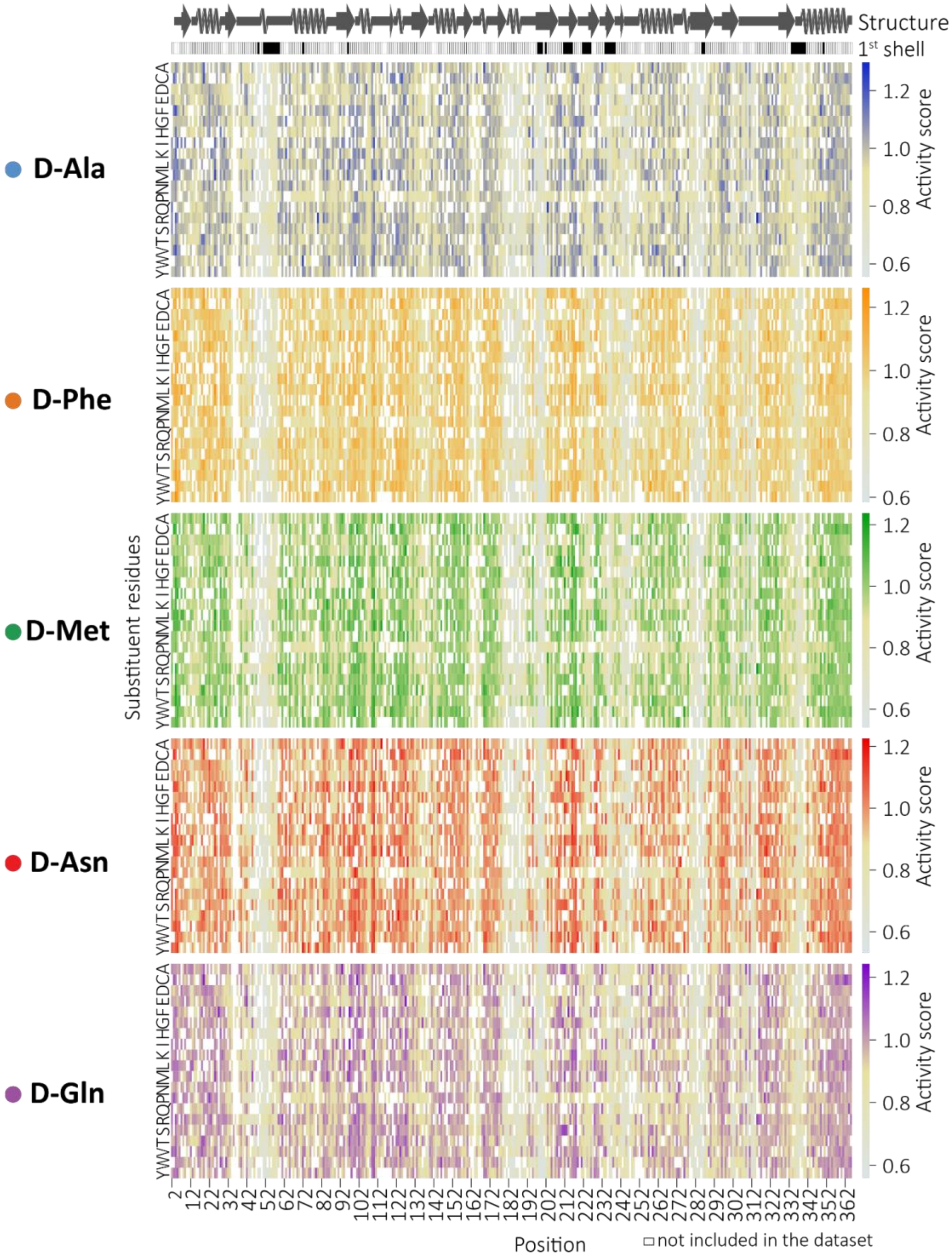

**Figure S4. DAOx single mutant activity scores across five substrates.**

Heatmaps displaying EP-Seq activity fitness scores for all single missense mutations screened with each of the five substrates. The intensity of the color on the heatmap reflects the relative impact of each mutation on enzymatic activity for the respective substrate.

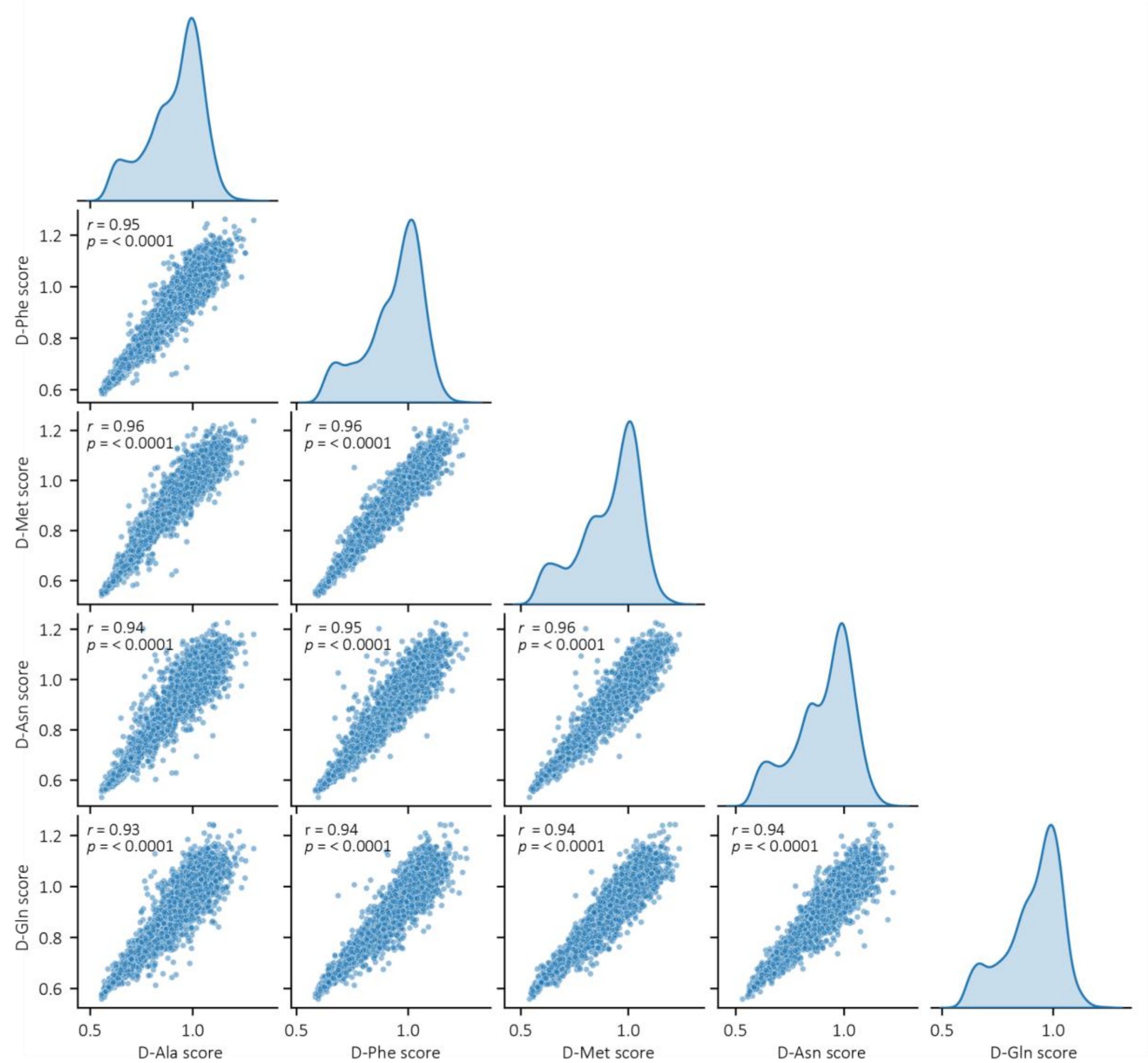

**Figure S5. Comparison of consensus DAOx activity fitness scores across five substrate screenings.**

Linear regression analyses (Pearson's *r*, two-tailed) were performed to compare the activity fitness scores from EP-Seq screenings using different DAOx substrates. Each plot represents the relationship between two distinct screening datasets. Pearson's r values indicate the degree of concordance between the datasets.

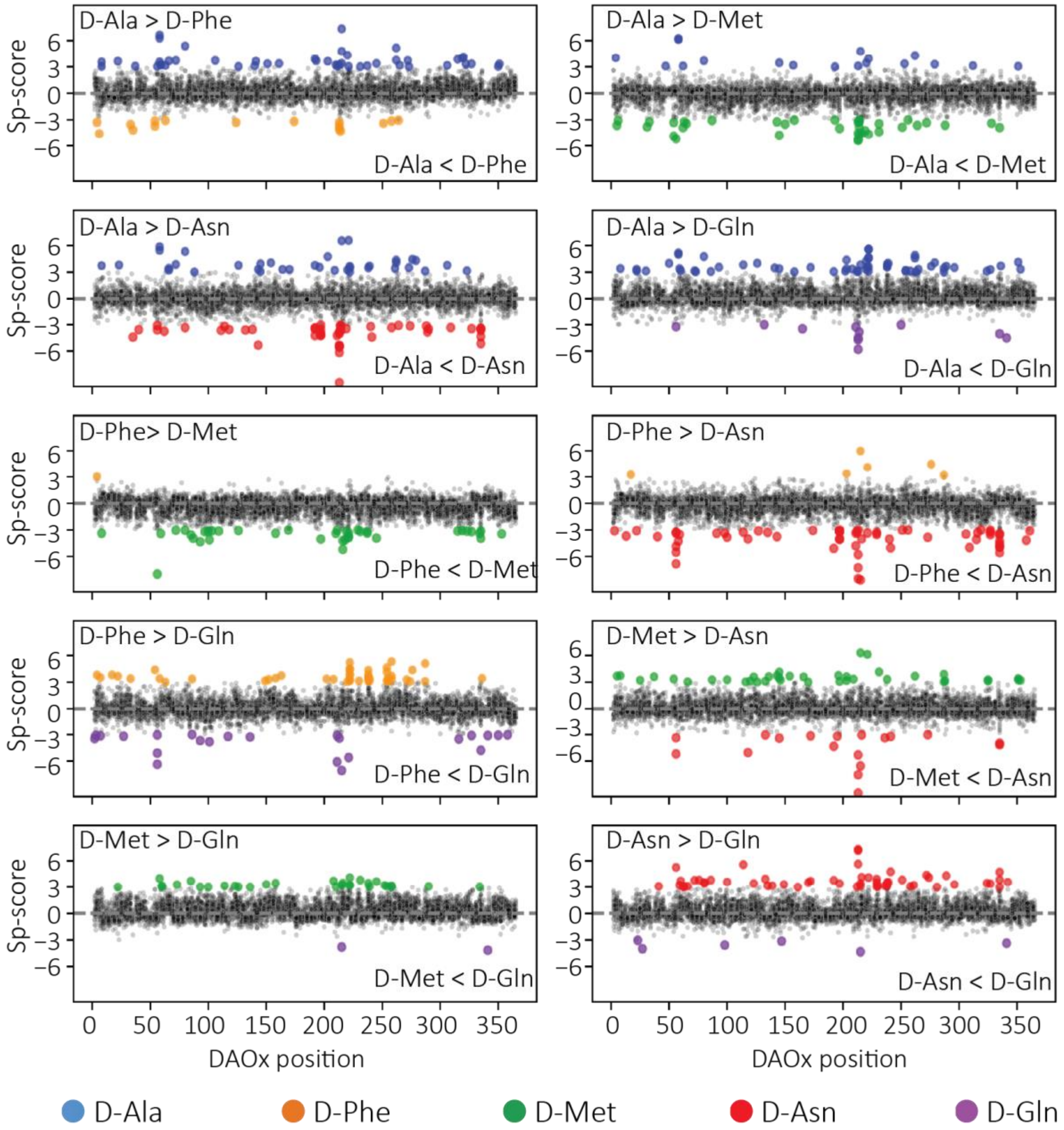

**Figure S6. Identification and localization of substrate-specific mutations in DAOx.**

Scatter plots of Sp-scores per position of DAOx for each variant enzyme and across all possible pairwise comparisons of EP-Seq screenings results performed with different substrates. Colored dots indicate variants with Sp-scores above 3 or below -3, representing substrate-specific mutations. Substrate-specific mutations are color-coded as follows: D-Ala-specific variants in blue, D-Phe-specific in orange, D-Met-specific in green, D-Asn-specific in red, and D-Gln-specific in violet. This color scheme is consistently used throughout the manuscript to denote substrate-specific mutations.

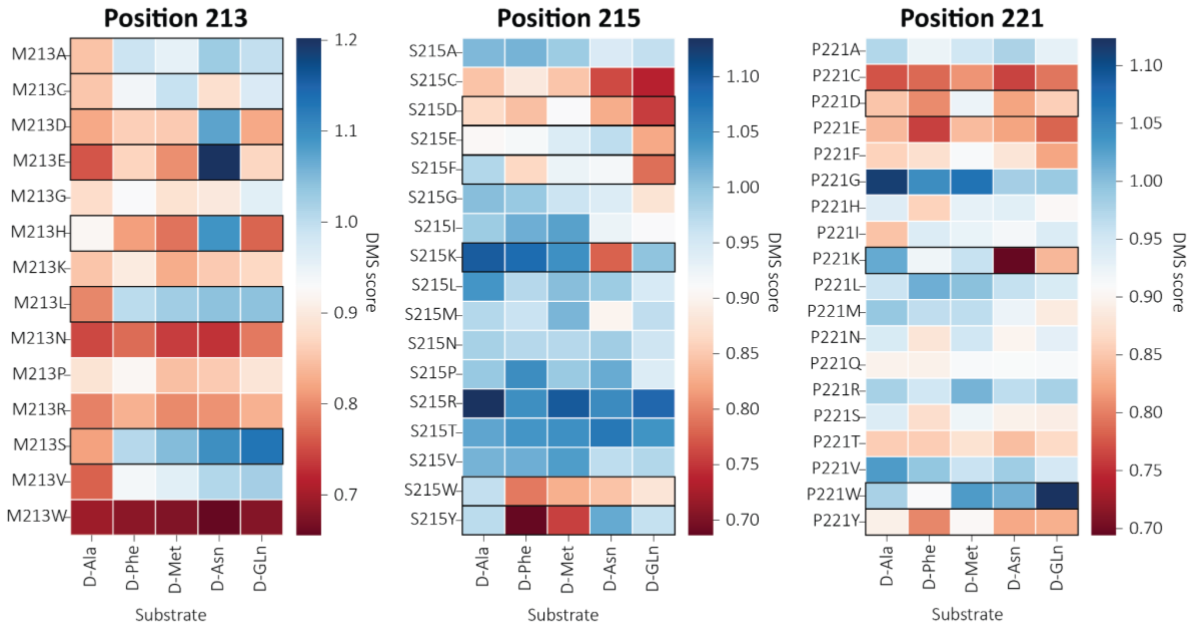

**Figure S7. DAOx positions 213, 215 and 221 play a general role in substrate specificity.**

Heatmap overview of EP-Seq scores for single-point mutants at positions 213, 215 and 221 tested across all five substrates (D-Ala, D-Phe, D-Met, D-Asn, and D-Gln), highlighting position-specific effects on substrate preference.

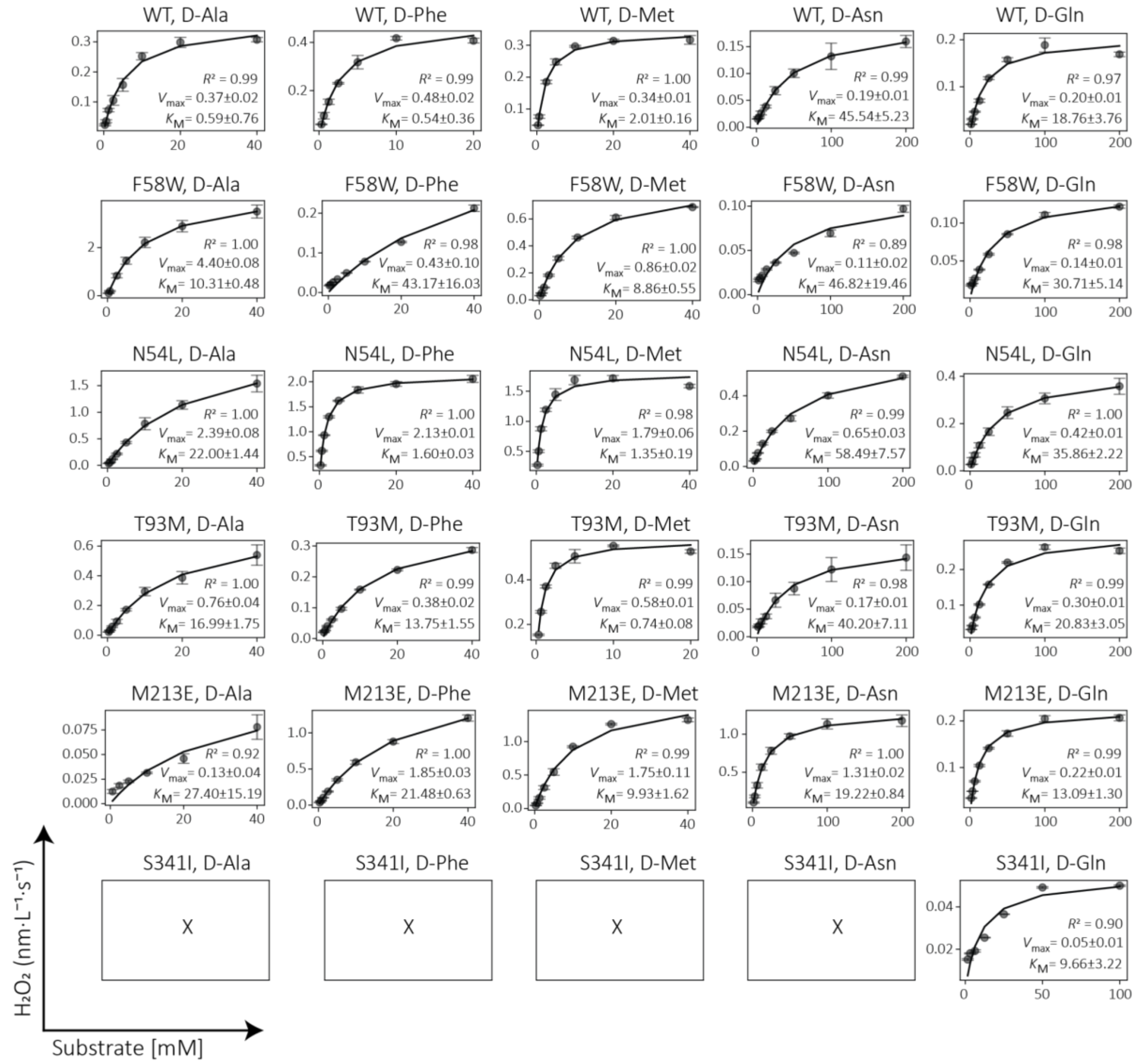

**Figure S8. Kinetic profiles of soluble WT and DAOx variants.**

Michaelis–Menten plots showing initial rates of $H_2O_2$ formation as a function of substrate concentration for wild-type (WT) and five DAOx variants. Soluble proteins were expressed in *E. coli* NiCo21(DE3) and purified using affinity and size exclusion chromatography to ensure high and comparable purity across all enzymes. Enzyme concentrations used for kinetic measurements were: 0.25 nM WT, 2 nM N54L, 2 nM F58W, 5 nM T93M, 5 nM S341I, and 1 nM M213E. X indicates no measurable activity, with signal below the detection limit of the assay.

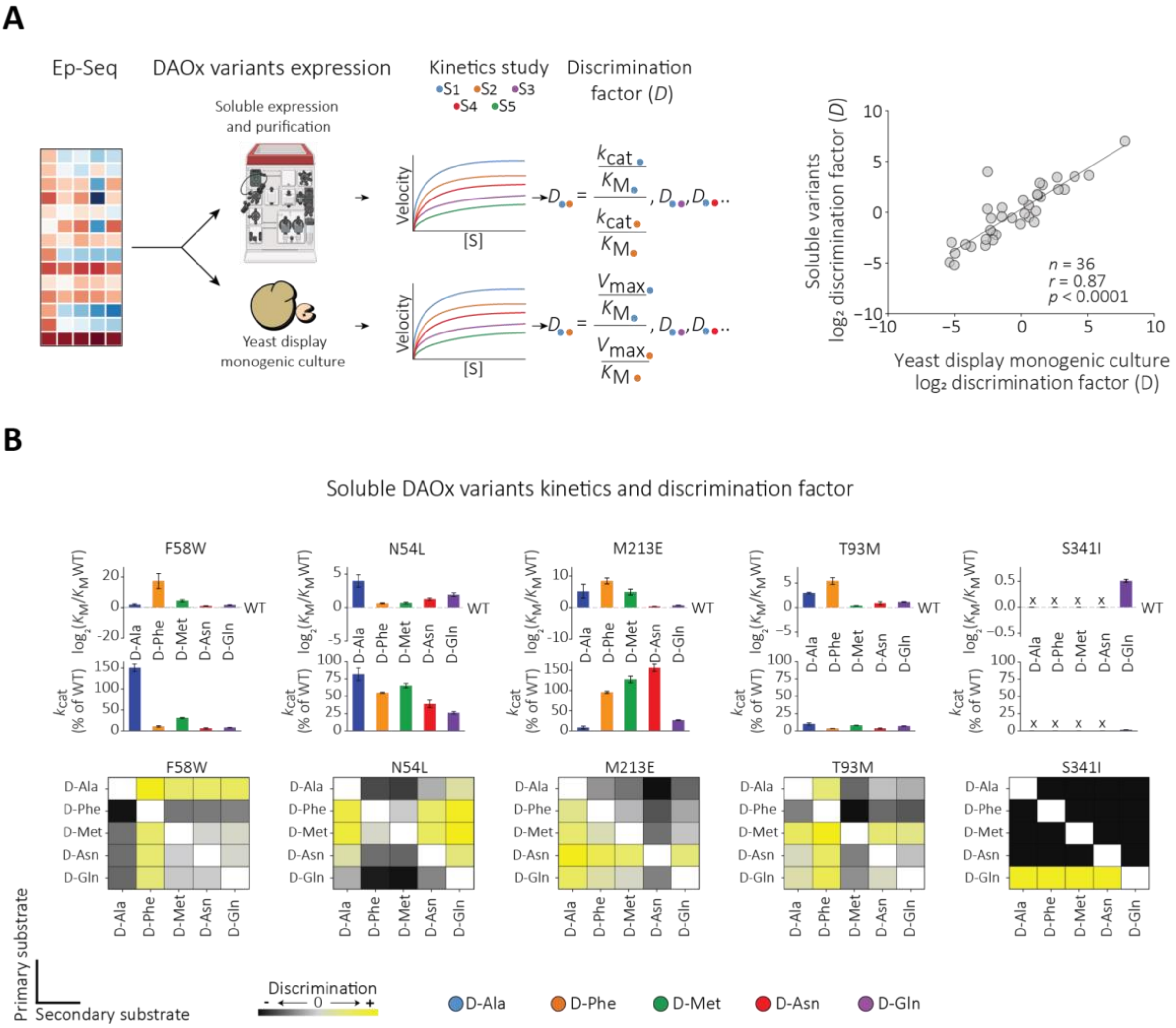

**Figure S9. Validation and Analysis of Soluble Variants' D-Factors.**

(**A**) Workflow illustrating the D-factor validation process: substrate-specific single mutants identified through EP-Seq were expressed both as monoclonal yeast display variants and as soluble proteins in *E. coli* (left). Catalytic efficiencies ($k_{cat}/K_M$) were determined for the soluble forms across all five substrates and compared with their yeast-displayed counterparts to assess consistency between formats (right). Dashed line: linear regression; *r*: Pearson correlation coefficients, n: number of datapoints, *p*: p-values two-tailed. (**B**)(Top) Catalytic profiles of selected variants expressed as soluble enzymes. Relative changes in $k_{cat}$ and $V_{max}$ values compared to the wild-type enzyme for each of the five substrates, illustrating how specificity is tuned through alterations in substrate affinity and catalytic turnover. (Bottom) Heatmap of substrate discrimination values between all pairwise combinations of primary (y-axis) and secondary (x-axis) substrates, quantifying the degree of specificity conferred by each mutation. X indicates no measurable activity, with signal below the detection limit of the assay.

### Supplementary Tables

**Table S1. Comparison of kinetic features between yeast-displayed and soluble WT-DAOx.**

Michaelis-Menten constant $K_M$ experimentally assayed for the yeast displayed and soluble form of the WT-DAOx. Coefficients of determination ($R^2$) for each of the fitted datasets are reported.

| Substrate | Soluble DAOx $K_M$ [mM] | $R^2$ | Surface displayed DAOx $K_M$ [mM] | $R^2$ |
|---|---|---|---|---|
| **D-Ala** | 6.965 ± 0.4003 | 0.9937 | 7.316 ± 0.2816 | 0.9973 |
| **D-Met** | 2.214 ± 0.1608 | 0.9850 | 2.039 ± 0.1455 | 0.9851 |
| **D-Phe** | 2.497 ± 0.1033 | 0.9950 | 1.507 ± 0.05914 | 0.9947 |
| **D-Gln** | 16.12 ± 0.8415 | 0.9961 | 14.42 ± 0.8434 | 0.9950 |
| **D-His** | 3.612 ± 0.1662 | 0.9942 | 3.406 ± 0.2003 | 0.9923 |
| **D-Thr** | 34.01 ± 2.745 | 0.9883 | 39.38 ± 1.548 | 0.9975 |
| **D-Asn** | 54.05 ± 4.683 | 0.9958 | 45.82 ± 1.488 | 0.9990 |
| **D-Ser** | 27.58 ± 2.755 | 0.9813 | 16.47 ± 0.6280 | 0.9967 |
| **D-Leu** | 5.021 ± 0.2623 | 0.9940 | 4.991 ± 0.1093 | 0.9990 |

**Table S2. Sorted cells, Illumina reads, and median fluorescence values per sorting gate**

| Experiment | Bin | # Illumina reads | # Sorted cells | Median fluorescence |
|---|---|---|---|---|
| D-Alanine screening Replicate 1 | 1 | 44100714 | 3951383 | 672.5 |
| | 2 | 37370660 | 425024 | 4580 |
| | 3 | 42886950 | 407658 | 17367 |
| | 4 | 41175573 | 485113 | 54275.125 |
| D-Alanine screening Replicate 2 | 1 | 49230428 | 7020636 | 504.5 |
| | 2 | 38379877 | 468261 | 4764.125 |
| | 3 | 37319082 | 625693 | 17491.625 |
| | 4 | 40512560 | 641907 | 52822.75 |
| D-Methionine screening Replicate 1 | 1 | 47604620 | 8290439 | 741 |
| | 2 | 36257935 | 1078896 | 5076 |
| | 3 | 33684162 | 1190286 | 26548 |
| | 4 | 34336748 | 1125873 | 71588 |
| D-Methionine screening Replicate 2 | 1 | 42408681 | 7565518 | 735 |
| | 2 | 35254873 | 971169 | 4986 |
| | 3 | 34252436 | 1032611 | 26473 |
| | 4 | 35238485 | 923507 | 71469 |
| D-Phenylalanine screening | 1 | 46910569 | 8343237 | 714 |
| | 2 | 28055538 | 1117236 | 5198 |

| | | | | |
|---|---|---|---|---|
| Replicate 1 | 3 | 27966355 | 1276152 | 28036 |
| | 4 | 25871098 | 1152820 | 75211 |
| D-Phenylalanine screening Replicate 2 | 1 | 37163633 | 6448964 | 690 |
| | 2 | 38860381 | 867477 | 5105 |
| | 3 | 32718506 | 870761 | 26350 |
| | 4 | 32567797 | 724807 | 70094 |
| D-Asparagine screening Replicate 1 | 1 | 42675792 | 6194903 | 744 |
| | 2 | 42086369 | 887066 | 4940 |
| | 3 | 31855534 | 920042 | 21483 |
| | 4 | 43640442 | 778517 | 70691 |
| D-Asparagine screening Replicate 2 | 1 | 35894362 | 6479013 | 773 |
| | 2 | 33143311 | 835295 | 4836 |
| | 3 | 27876393 | 939669 | 22120 |
| | 4 | 30558626 | 840663 | 71013 |
| D-Glutamine screening Replicate 1 | 1 | 41006142 | 6663201 | 484 |
| | 2 | 38484786 | 912438 | 2650 |
| | 3 | 43725266 | 925690 | 5522 |
| | 4 | 43218258 | 950405 | 17090 |
| D-Glutamine screening | 1 | 28002452 | 5918410 | 485 |
| | 2 | 40842045 | 872992 | 2664 |

| | | | | |
|---|---|---|---|---|
| Replicate 2 | 3 | 30320723 | 897982 | 5534 |
| | 4 | 32201970 | 947327 | 17639 |

#### Table S3. Linear regression correlation and fitness metrics per subpopulation in EP-Seq screening

Pearson correlation coefficient (PCCR) *r* of the linear regression between experimental replicates in **Fig. 2**. Details on the composition of the single mutant variants per each EP-Seq screen and substrate. Specifically, number of missense variants, percentage of total, median score; number of nonsense variants and average activity score with standard deviation; number of synonymous variants and average activity score with standard deviation.

| | |
|---|---|
| **D-Alanine** | |
| PCC (*r*) | 0.93 |
| # missense variants # (% tot, median score) | 6377 (92.2%, -0.33) |
| # nonsense variants (mean score ± SD) | 332 (-0.60±0.11) |
| # synonymous variants (mean score ± SD) | 298 (-0.000565±0.064) |
| **D-Phenylalanine** | |
| PCC (*r*) | 0.96 |
| # missense variants # (% tot, median score) | 6400 (92.5%, -0.28) |
| # nonsense variants (mean score ± SD) | 333 (-0.53±0.09) |
| # synonymous variants (mean score ± SD) | 300 (-0.000546±0.054) |
| **D-Methionine** | |
| PCC (*r*) | 0.96 |
| # missense variants # (% tot, median score) | 6414 (92.7%, -0.28) |
| # nonsense variants (mean score ± SD) | 333 (-0.63±0.11) |
| # synonymous variants (mean score ± SD) | 301 (-0.00368±0.052) |
| **D-Glutamine** | |
| PCC (r) | 0.94 |
| # missense variants # (% tot, median score) | 6426 (92.9%, -0.35) |
| # nonsense variants (mean score ± SD) | 332 (-0.55±0.10) |
| # synonymous variants (mean score ± SD) | 301 (-0.00232±0.051) |
| **D-Asparagine** | |
| PCC (*r*) | 0.93 |
| # missense variants # (% tot, median score) | 6379 (92.2%, -0.34) |
| # nonsense variants (mean score ± SD) | 332 (-0.61±0.11) |
| # synonymous variants (mean score ± SD) | 302 (-0.0007±0.061) |
| **All datasets (variants shared by all datasets)** | |
| # missense variants # (% tot, mean score) | 5821 (84.2%, -0.31±0.05) |
| # nonsense variants (mean score ± SD) | 316 (-0.58±0.05) |
| # synonymous variants (mean score ± SD) | 281 (-0.0026±0.0350) |

#### Table S4. Most selective DAOx variants per substrate identified by EP-Seq

For each variant, the table reports the distance of the mutation from the catalytic site, the identities of the primary and secondary substrates, the measured activity scores on each substrate, and the Sp-score between the primary and secondary substrates.

| | | Substrate | | Activity score | | | | | |
|---|---|---|---|---|---|---|---|---|---|
| DAOx variant | Distance to catalytic site (Å) | Primary | Secondary | D-Ala | D-Phe | D-Met | D-Asn | D-Gln | Sp-score |
| **S215Y** | 7.28 | **D-Ala** | D-Phe | 0.97 | 0.69 | 0.76 | 1.03 | 0.96 | 7.33 |
| **F58Y** | 2.21 | **D-Ala** | D-Phe | 0.92 | 0.66 | 0.64 | 0.63 | 0.65 | 6.65 |
| **P221K** | 7.10 | **D-Ala** | D-Asn | 1.02 | 0.92 | 0.96 | 0.69 | 0.84 | 6.57 |
| **S215K** | 7.28 | **D-Ala** | D-Asn | 1.10 | 1.08 | 1.05 | 0.78 | 1.00 | 6.54 |
| **F58W** | 2.21 | **D-Ala** | D-Phe | 0.90 | 0.66 | 0.62 | 0.63 | 0.64 | 6.30 |
| **F58Y** | 2.21 | **D-Ala** | D-Met | 0.92 | 0.66 | 0.64 | 0.63 | 0.65 | 6.25 |
| **F58W** | 2.21 | **D-Ala** | D-Met | 0.90 | 0.66 | 0.62 | 0.63 | 0.64 | 6.13 |
| **F58Y** | 2.21 | **D-Ala** | D-Asn | 0.92 | 0.66 | 0.64 | 0.63 | 0.65 | 5.86 |
| **A222M** | 6.74 | **D-Ala** | D-Gln | 0.99 | 0.91 | 0.85 | 0.80 | 0.70 | 5.60 |
| **A222N** | 6.74 | **D-Ala** | D-Gln | 1.13 | 1.09 | 0.95 | 0.95 | 0.84 | 5.56 |
| **M213E** | 2.51 | **D-Asn** | D-Met | 0.76 | 0.87 | 0.80 | 1.20 | 0.87 | 9.61 |
| **M213E** | 2.51 | **D-Asn** | D-Ala | 0.76 | 0.87 | 0.80 | 1.20 | 0.87 | 9.57 |
| **S215Y** | 7.28 | **D-Asn** | D-Phe | 0.97 | 0.69 | 0.76 | 1.03 | 0.96 | 8.66 |
| **M213E** | 2.51 | **D-Asn** | D-Phe | 0.76 | 0.87 | 0.80 | 1.20 | 0.87 | 8.51 |
| **M213H** | 2.51 | **D-Asn** | D-Met | 0.92 | 0.81 | 0.78 | 1.09 | 0.77 | 7.54 |
| **M213E** | 2.51 | **D-Asn** | D-Gln | 0.76 | 0.87 | 0.80 | 1.20 | 0.87 | 7.32 |
| **M213H** | 2.51 | **D-Asn** | D-Phe | 0.92 | 0.81 | 0.78 | 1.09 | 0.77 | 7.28 |
| **M213H** | 2.51 | **D-Asn** | D-Gln | 0.92 | 0.81 | 0.78 | 1.09 | 0.77 | 7.18 |
| **T56H** | 3.47 | **D-Asn** | D-Phe | 0.81 | 0.71 | 0.76 | 0.97 | 0.74 | 6.84 |
| **S215Y** | 7.28 | **D-Asn** | D-Met | 0.97 | 0.69 | 0.76 | 1.03 | 0.96 | 6.55 |
| **T56M** | 3.47 | **D-Met** | D-Phe | 0.83 | 0.76 | 1.05 | 0.93 | 1.01 | 7.99 |
| **S215K** | 7.28 | **D-Met** | D-Asn | 1.10 | 1.08 | 1.05 | 0.78 | 1.00 | 6.32 |
| **P221K** | 7.10 | **D-Met** | D-Asn | 1.02 | 0.92 | 0.96 | 0.69 | 0.84 | 6.13 |
| **M213S** | 2.51 | **D-Met** | D-Ala | 0.82 | 1.01 | 1.05 | 1.10 | 1.13 | 5.39 |
| **M213L** | 2.51 | **D-Met** | D-Ala | 0.80 | 1.00 | 1.02 | 1.04 | 1.04 | 5.28 |
| **S216G** | 10.84 | **D-Met** | D-Phe | 1.01 | 1.03 | 1.21 | 1.11 | 1.04 | 5.20 |

| | | | | | | | | | |
|---|---|---|---|---|---|---|---|---|---|
| **T56M** | 3.47 | **D-Met** | D-Ala | 0.83 | 0.76 | 1.05 | 0.93 | 1.01 | 5.18 |
| **N54I** | 2.58 | **D-Met** | D-Ala | 0.69 | 0.86 | 0.90 | 0.75 | 0.77 | 4.89 |
| **Y145W** | 17.59 | **D-Met** | D-Ala | 0.92 | 1.06 | 1.13 | 0.98 | 1.04 | 4.79 |
| **D214F** | 6.73 | **D-Met** | D-Ala | 0.98 | 1.09 | 1.18 | 1.11 | 1.18 | 4.72 |
| **S215K** | 7.28 | **D-Phe** | D-Asn | 1.10 | 1.08 | 1.05 | 0.78 | 1.00 | 5.99 |
| **H258W** | 11.57 | **D-Phe** | D-Gln | 1.06 | 1.23 | 1.10 | 1.10 | 0.97 | 5.32 |
| **A222N** | 6.74 | **D-Phe** | D-Gln | 1.13 | 1.09 | 0.95 | 0.95 | 0.84 | 5.22 |
| **A287H** | 8.78 | **D-Phe** | D-Gln | 1.00 | 1.07 | 0.95 | 0.88 | 0.82 | 5.10 |
| **R254W** | 10.28 | **D-Phe** | D-Gln | 1.03 | 1.15 | 1.07 | 1.02 | 0.93 | 4.63 |
| **R6A** | 32.00 | **D-Phe** | D-Ala | 0.86 | 1.09 | 0.99 | 0.95 | 1.02 | 4.59 |
| **A222H** | 6.74 | **D-Phe** | D-Gln | 0.91 | 0.98 | 0.91 | 0.81 | 0.76 | 4.52 |
| **E276F** | 17.89 | **D-Phe** | D-Asn | 1.10 | 1.20 | 1.06 | 0.96 | 0.98 | 4.47 |
| **A222M** | 6.74 | **D-Phe** | D-Gln | 0.99 | 0.91 | 0.85 | 0.80 | 0.70 | 4.41 |
| **N54L** | 2.58 | **D-Phe** | D-Gln | 0.70 | 0.89 | 0.81 | 0.74 | 0.67 | 4.39 |
| **S215Y** | 7.28 | **D-Gln** | D-Phe | 0.97 | 0.69 | 0.76 | 1.03 | 0.96 | 7.06 |
| **T56M** | 3.47 | **D-Gln** | D-Phe | 0.83 | 0.76 | 1.05 | 0.93 | 1.01 | 6.36 |
| **C211S** | 5.99 | **D-Gln** | D-Phe | 1.00 | 0.90 | 0.99 | 1.07 | 1.14 | 6.08 |
| **M213S** | 2.51 | **D-Gln** | D-Ala | 0.82 | 1.01 | 1.05 | 1.10 | 1.13 | 5.81 |
| **P221W** | 7.10 | **D-Gln** | D-Phe | 0.98 | 0.91 | 1.03 | 1.01 | 1.12 | 5.60 |
| **T56Q** | 3.47 | **D-Gln** | D-Phe | 0.90 | 0.83 | 0.90 | 1.03 | 1.03 | 5.09 |
| **S335K** | 2.07 | **D-Gln** | D-Phe | 0.77 | 0.81 | 0.94 | 0.93 | 0.99 | 4.78 |
| **M213V** | 2.51 | **D-Gln** | D-Ala | 0.77 | 0.94 | 0.96 | 1.01 | 1.02 | 4.73 |
| **M213L** | 2.51 | **D-Gln** | D-Ala | 0.80 | 1.00 | 1.02 | 1.04 | 1.04 | 4.51 |
| **S341I** | 10.55 | **D-Gln** | D-Ala | 0.81 | 0.95 | 0.83 | 0.88 | 1.06 | 4.51 |

#### Table S5. Kinetic profiles of yeast-displayed WT and DAOx variants.

Kinetic parameters ($K_M$ and $V_{max}$) for each yeast-displayed DAOx variant tested individually against the five substrates used in this study. "X" indicates values that were not determined due to either the absence of detectable activity or unreliable fitting of the kinetic data. WT-DAOx was measured in parallel with each variant. $V_{max}$ values for each variant were normalized to their respective expression levels, as determined by immunostaining, prior to being reported in the table for comparison. The WT reference for each variant is indicated by the same color code in the table.

| DAOx variants (yeast display) | Substrate | | | | | | | | | |
|---|---|---|---|---|---|---|---|---|---|---|
| | D-Alanine | | D-Phenylalanine | | D-Methionine | | D-Asparagine | | D-Glutamine | |
| | $K_M$ [mM] | $V_{max}$ (nM/sec) | $K_M$ [mM] | $V_{max}$ (nM/sec) | $K_M$ [mM] | $V_{max}$ (nM/sec) | $K_M$ [mM] | $V_{max}$ (nM/sec) | $K_M$ [mM] | $V_{max}$ (nM/sec) |
| **WT_1** | 2.31±0.72 | 1.94±0.23 | 0.45±0.07 | 1.99±0.14 | 0.49±0.01 | 2.04±0.03 | 9.66±0.35 | 0.93±0.03 | 7.94±0.32 | 0.75±0.01 |
| **N54I** | 16.05±1.43 | 0.54±0.03 | 0.85±0.04 | 0.67±0.0 | 0.49±0.02 | 0.75±0.01 | 21.05±2.62 | 0.21±0.0 | 10.6±0.21 | 0.23±0.0 |
| **N54L** | 22.24±0.91 | 0.76±0.0 | 0.83±0.02 | 0.65±0.01 | 0.91±0.25 | 0.56±0.0 | 26.04±0.58 | 0.16±0.01 | 25.11±0.86 | 0.14±0.0 |
| **T56H** | 7.15±0.25 | 0.7±0.0 | 16.03±0.22 | 1.34±0.01 | 3.59±0.16 | 0.95±0.02 | 22.77±0.68 | 0.76±0.02 | 23.93±1.37 | 0.24±0.01 |
| **T56M** | 12.68±0.68 | 1.1±0.02 | 15.18±0.34 | 1.2±0.04 | 0.53±0.01 | 1.09±0.01 | 31.44±8.38 | 0.64±0.0 | 7.81±0.8 | 0.51±0.02 |
| **T56Q** | 11.83±2.53 | 1.22±0.04 | 7.1±0.57 | 1.37±0.03 | 1.83±0.12 | 1.22±0.04 | 10.27±1.97 | 0.61±0.05 | 8.29±0.63 | 0.55±0.01 |
| **F58W** | 7.26±0.05 | 1.44±0.01 | 65.79±27.12 | 0.3±0.12 | 5.5±0.13 | 0.33±0.0 | 47.88±0.57 | 0.03±0.0 | 20.56±1.07 | 0.08±0.0 |
| **T93M** | 10.8±0.26 | 0.97±0.01 | 17.01±1.03 | 0.83±0.03 | 0.37±0.02 | 0.71±0.0 | 32.21±10.35 | 0.22±0.01 | 19.62±0.95 | 0.38±0.01 |
| **I197S** | 1.96±0.09 | 0.52±0.02 | 2.16±0.12 | 1.23±0.06 | 0.69±0.05 | 0.89±0.03 | 14.71±0.1 | 0.74±0.01 | 10.97±0.9 | 0.45±0.02 |
| **M213D** | 55.9±4.75 | 0.35±0.02 | 28.24±0.6 | 1.09±0.03 | 4.37±0.03 | 0.61±0.0 | 8.36±1.69 | 0.53±0.01 | 5.02±0.55 | 0.07±0.0 |
| **M213E** | X | X | 10.34±3.64 | 1.07±0.2 | 9.72±2.84 | 0.99±0.18 | 8.1±1.88 | 0.61±0.12 | 12.37±0.59 | 0.14±0.0 |
| **M213H** | 10.89±2.38 | 1.57±0.08 | 5.78±0.11 | 1.29±0.01 | 4.93±0.22 | 1.05±0.0 | 11.38±0.38 | 1.0±0.01 | 11.69±0.64 | 0.22±0.01 |
| **S215Y** | 2.08±0.12 | 0.89±0.0 | 14.84±0.05 | 0.71±0.0 | 2.07±0.11 | 0.59±0.0 | 6.39±0.32 | 0.58±0.01 | 8.53±0.24 | 0.34±0.0 |
| **A222M** | 4.44±0.26 | 1.05±0.08 | 1.5±0.21 | 1.02±0.1 | 2.43±0.04 | 0.86±0.03 | 21.11±0.54 | 0.27±0.01 | 26.91±0.41 | 0.17±0.0 |
| **H258W** | 6.51±0.5 | 1.72±0.07 | 0.55±0.01 | 1.42±0.02 | 1.34±0.05 | 1.59±0.01 | 11.15±0.32 | 0.46±0.01 | 18.08±0.07 | 0.45±0.0 |
| **E276F** | 4.48±0.81 | 2.47±0.16 | 0.77±0.1 | 2.36±0.1 | 0.91±0.1 | 2.43±0.06 | 18.14±1.65 | 1.0±0.0 | 13.9±1.33 | 0.81±0.02 |
| **S335K** | X | X | 6.93±0.03 | 0.39±0.0 | 1.47±0.0 | 0.71±0.0 | 37.57±0.65 | 0.57±0.0 | 10.41±0.27 | 0.43±0.0 |
| **S341I** | X | X | X | X | X | X | X | X | 11.17±0.87 | 0.06±0.0 |
| **WT_2** | 2.4±0.07 | 2.1±0.06 | 0.79±0.1 | 2.45±0.06 | 0.65±0.0 | 2.08±0.01 | 19.6±0.8 | 1.06±0.08 | 16.2±2.41 | 0.85±0.03 |
| **Y145A** | 3.0±0.08 | 1.45±0.08 | 0.79±0.04 | 1.61±0.02 | 0.94±0.03 | 1.56±0.08 | 23.4±3.66 | 0.63±0.07 | 17.27±0.15 | 0.57±0.01 |
| **Y145W** | 3.49±0.01 | 1.71±0.02 | 0.78±0.01 | 1.62±0.01 | 0.98±0.02 | 1.66±0.06 | 25.01±0.39 | 0.76±0.0 | 18.42±0.05 | 0.56±0.01 |

| | | | | | | | | | | |
|---|---|---|---|---|---|---|---|---|---|---|
| **Q192C** | 3.55±0.23 | 1.11±0.02 | 0.87±0.07 | 1.13±0.02 | 1.02±0.03 | 1.23±0.0 | 24.61±0.5 | 0.58±0.01 | 16.85±0.48 | 0.44±0.0 |
| **Q192P** | 2.56±0.0 | 0.94±0.05 | 0.7±0.02 | 0.89±0.01 | 0.79±0.01 | 0.98±0.01 | 22.04±1.34 | 0.47±0.01 | 17.02±0.07 | 0.39±0.0 |
| **S216G** | 3.62±0.06 | 1.93±0.05 | 0.37±0.01 | 1.73±0.02 | 0.6±0.01 | 1.94±0.01 | 20.14±5.19 | 0.96±0.05 | 13.25±0.87 | 0.78±0.01 |
| **S216H** | 2.08±0.16 | 2.12±0.19 | 1.08±0.04 | 2.25±0.04 | 1.0±0.02 | 2.05±0.0 | 21.06±1.26 | 0.92±0.0 | 18.52±0.3 | 0.65±0.02 |
| **S220H** | 3.0±1.03 | 1.67±0.07 | 0.31±0.04 | 1.53±0.02 | 0.65±0.02 | 1.45±0.0 | 21.22±8.72 | 0.85±0.38 | 18.18±0.3 | 0.63±0.0 |
| **S220V** | 2.82±0.04 | 1.91±0.03 | 0.45±0.02 | 1.76±0.03 | 0.84±0.1 | 1.7±0.06 | 19.35±1.1 | 0.83±0.07 | 18.34±1.11 | 0.59±0.02 |
| **R254C** | 3.3±0.0 | 1.45±0.0 | 0.47±0.02 | 1.5±0.01 | 1.52±0.49 | 1.4±0.13 | 23.3±0.05 | 0.55±0.01 | 21.38±0.12 | 0.39±0.0 |
| **R254D** | 5.62±0.53 | 1.07±0.02 | 1.02±0.04 | 1.11±0.03 | 1.67±0.07 | 1.1±0.0 | 35.04±0.89 | 0.39±0.01 | 19.99±0.66 | 0.28±0.0 |
| **H258R** | 5.96±0.7 | 1.32±0.02 | 0.63±0.01 | 1.15±0.02 | 1.89±0.09 | 1.2±0.02 | 41.29±0.7 | 0.43±0.1 | 23.4±0.85 | 0.25±0.0 |
| **L262A** | 5.66±0.42 | 1.04±0.01 | 2.79±0.02 | 1.04±0.06 | 2.17±0.4 | 0.99±0.05 | 57.29±0.52 | 0.33±0.0 | 26.56±0.07 | 0.24±0.0 |
| **L262V** | 2.86±0.06 | 1.24±0.08 | 2.52±0.02 | 1.53±0.06 | 1.74±0.08 | 1.47±0.02 | 34.08±1.93 | 0.48±0.0 | 33.16±1.28 | 0.5±0.02 |
| **M213E S215Y** | 30.37±2.94 | 0.11±0.01 | 26.84±4.73 | 0.12±0.02 | 10.6±0.72 | 0.07±0.0 | 4.31±0.91 | 0.05±0.0 | 14.89±0.66 | 0.2±0.0 |
| **F58W S215Y** | 5.48±0.15 | 0.89±0.01 | X | X | 6.99±0.06 | 0.16±0.0 | 70.19±7.76 | 0.13±0.01 | 20.02±1.92 | 0.13±0.0 |
| **F58W M213E** | X | X | X | X | 11.81±0.27 | 0.14±0.0 | 29.92±0.63 | 0.13±0.0 | 17.36±3.88 | 0.08±0.0 |
| **F58W A222M** | 12.09±0.63 | 0.81±0.05 | X | X | 12.7±0.9 | 0.07±0.0 | X | X | 16.82±0.32 | 0.03±0.0 |
| **N54L H258W** | 23.46±3.1 | 0.37±0.02 | 0.83±0.02 | 0.63±0.04 | 1.37±0.13 | 0.29±0.0 | 56.58±7.15 | 0.11±0.0 | 24.05±1.93 | 0.1±0.0 |
| **T56H T93M** | 4.78±0.09 | 0.66±0.02 | 20.67±0.71 | 1.11±0.0 | 6.64±0.51 | 0.8±0.01 | 77.6±70.09 | 0.49±0.34 | 30.07±1.53 | 0.18±0.0 |

1. Vanella, R. *et al.* Understanding activity-stability tradeoffs in biocatalysts by enzyme proximity sequencing. *Nature Communications* **15**, 1807 (2023).
2. Gietz, R. D. & Woods, R. A. Transformation of yeast by lithium acetate/single-stranded carrier DNA/polyethylene glycol method. *Methods Enzymol.* **350**, 87–96 (2002).
3. Bushnell, B. BBMap: A Fast, Accurate, Splice-Aware Aligner. *https://www.semanticscholar.org › paper › BBMap:-A-Fas...https://www.semanticscholar.org › paper › BBMap:-A-Fas...* (2014).
